# Evolution of regulatory chromatin contacts: insights from duplicated genes

**DOI:** 10.64898/2026.07.22.739131

**Authors:** Victor Lefebvre, Sylvain Foissac, Sarah Djebali, Anamaria Necsulea

## Abstract

In multicellular organisms, gene expression is controlled by numerous cis-regulatory elements that can be located far away from their target genes on the linear genome. Interactions between gene promoters and distant regulatory elements take place through chromatin contacts or loops. While other aspects of gene expression regulation have been well studied from an evolutionary perspective, regulatory chromatin contact evolution remains largely unexplored, due to a lack of comparable data across species. Here, we study the evolution of regulatory chromatin contacts by focusing on duplicated genes in the mouse genome. We use an extensive collection of high resolution promoter-centered Hi-C data to define promoter-enhancer chromatin contacts for 1,420 pairs of duplicated genes and to study their evolutionary divergence, in conjunction with the evolutionary divergence of their expression patterns. We show that chromatin contacts evolve faster than expression patterns, as previously observed for other gene regulation mechanisms. We show that duplicated gene localisation in cis or in trans is strongly associated with chromatin contact divergence. We find a significant correlation between expression divergence and chromatin contact divergence across duplicated gene pairs. Our results highlight the complex evolutionary dynamics of regulatory chromatin contacts and the association between chromatin contact evolution and gene expression evolution, across a broad time scale.

## Introduction

In multicellular organisms, the regulation of gene expression is complex. At the transcriptional level, gene expression is controlled through interactions between *cis*-acting DNA elements (*i.e.* proximal promoter elements and more distant enhancers and silencers) and *trans*-acting factors (*i.e.* proteins and RNA molecules that bind to *cis*-acting elements). The transcriptional control of gene expression is tightly associated with the conformation of the chromatin. In particular, the three-dimensional organization of the chromatin is known to play an important role in gene expression regulation, as distant *cis*-regulatory elements (CREs) and gene promoters are brought into physical proximity in the nucleus through the formation of chromatin loops (Sexton, Bantignies, and Cavalli 2009; Montavon and Duboule 2012).

The evolution of regulatory mechanisms has been a major research focus in evolutionary biology since the seminal work of King and Wilson, who proposed that regulatory differences may account for many or even most phenotypic differences between species (King and Wilson 1975). Thus, several facets of gene expression regulation have been compared across species, bringing insights into the evolutionary dynamics of regulatory mechanisms. For example, the evolution of transcription factor binding was studied through comparative analyses of chromatin immunoprecipitation followed by sequencing (ChIP-seq) data, in various taxonomic groups (Schmidt et al. 2010; Schmidt et al. 2012; Paris et al. 2013; Ballester et al. 2014; Denas et al. 2015; Muiño et al. 2016). The evolution of gene promoters and distant enhancer elements was likewise studied through comparative ChIP-seq analyses, which targeted histone modifications that are characteristic of these classes of CREs (Villar et al. 2015; Berthelot et al. 2018). These studies revealed that there is extensive variation in the rates and patterns of evolution for different types of regulatory mechanisms. Notably, distal CREs (in particular enhancer sequences, activities or individual transcription factor binding locations) were shown to evolve rapidly, while proximal CREs (gene promoters), transcription factor sequences and binding motifs tend to be highly conserved across species.

Compared to other transcriptional regulatory mechanisms, the involvement of the three-dimensional (3D) chromatin organization in gene expression control has been less well studied from an evolutionary perspective. The 3D chromatin organization can be investigated with genome-wide chromosome conformation capture techniques such as Hi-C (Lieberman-Aiden et al. 2009), Micro-C (Hsieh et al. 2015) and related approaches. These techniques have revealed large-scale organizational features of the genome, including the existence of topologically-associating domains (Nora et al. 2012; Dixon et al. 2012), which are megabase-scale genomic regions inside which chromatin contacts occur preferentially. The evolution of these higher-order structures was investigated in detail, through comparative analyses of Hi-C data (Dixon et al. 2012; Eres et al. 2019; Gilbertson et al. 2022; Okhovat et al. 2023; Patalano et al. 2025). However, the evolution of fine-scale chromatin contacts between gene promoters and distant CREs is still largely unexplored. The scarcity of large-scale comparative studies of promoter-CRE chromatin contacts can likely be attributed to technical challenges. Indeed, while Hi-C can now be readily applied to a wide range of species, this technique is well suited for the characterization of higher-order chromatin organization features but lacks the power to identify fine-scale chromatin contacts. Several derived techniques, such as capture Hi-C (Schoenfelder et al. 2015; Mifsud et al. 2015) and capture Micro-C (Hua et al. 2021), were specifically designed to increase detection power for fine-scale chromatin contacts, by targeting a pre-defined set of regions of interest. In particular, promoter capture Hi-C (PCHi-C) can identify promoter-centered chromatin contacts with high sensitivity and resolution (Schoenfelder et al. 2015; Mifsud et al. 2015). Publicly available PCHi-C datasets for human and mouse were exploited in a comparative analysis of fine-scale promoter-CRE contacts (Laverré, Tannier, and Necsulea 2022). However, this study was limited by the availability of PCHi-C data for only two species, which greatly restricted the scope of the evolutionary analysis. Moreover, publicly available PCHi-C data were not necessarily derived from comparable biological conditions for the two species, meaning that the extent of evolutionary conservation could not be accurately estimated. To perform a broad evolutionary comparison of promoter-CRE chromatin contacts, generating comparable, high-resolution data across multiple species is needed. However, implementing promoter capture Hi-C protocols remains both technically challenging and costly.

Here, we aimed to study the evolution of promoter-enhancer contacts for duplicated genes, in the mouse genome. Duplication events are a widespread feature of genome evolution. In particular, vertebrate genomes harbour numerous instances of duplicated genes (also called paralogous genes), that are derived either from the duplication of restricted genomic segments or from the two rounds of whole-genome duplications that occurred in the vertebrate ancestor (Ohno 1970; Dehal and Boore 2005). Genome-wide analyses revealed that these gene duplications span a wide range of divergence times, from very recent events to events that occurred several hundreds of million years ago (Mya) (Lander et al. 2001; Mouse Genome Sequencing Consortium 2002; Friedman and Hughes 2004). Thus, comparing promoter-CRE chromatin contacts between duplicated genes can provide insights into their evolutionary dynamics across broad evolutionary timescales, while using chromatin conformation capture data generated for a single species.

The evolution of duplicated gene expression patterns has been extensively studied (Li, Yang, and Gu 2005; Kryuchkova-Mostacci and Robinson-Rechavi 2016; Lan and Pritchard 2016; Guschanski, Warnefors, and Kaessmann 2017). The regulatory characteristics of duplicated genes and the regulatory changes that may underlie expression divergence between paralogous gene copies have likewise attracted considerable attention. In particular, it was recently shown that duplicated gene promoter sequences evolve rapidly compared to orthologous gene promoters (Fraimovitch and Hagai 2023), which is consistent with the patterns observed for duplicated gene expression (Kryuchkova-Mostacci and Robinson-Rechavi 2016; Lan and Pritchard 2016; Guschanski, Warnefors, and Kaessmann 2017). The three-dimensional chromatin environment of duplicated genes was likewise scrutinized (Ibn-Salem, Muro, and Andrade-Navarro 2017). This study showed that paralogous gene pairs tend to be co-localized within the same topologically-associating domains and that they have high chromatin contact frequencies even when situated in different domains (Ibn-Salem, Muro, and Andrade-Navarro 2017). These observations suggest that the three-dimensional chromatin organization of duplicated genes may be involved in their co-regulation. This raises the question of how the divergence of the three-dimensional chromatin environment of paralogous genes may be associated with the divergence of their expression patterns.

Here, we address this question by studying the evolution of regulatory chromatin contacts and the evolution of expression patterns, for mouse duplicated genes. We focus on promoter-enhancer chromatin contacts, defined with high resolution Promoter Capture Hi-C data (Schoenfelder et al. 2015; Mifsud et al. 2015). Our results reveal that expression profiles and chromatin contact profiles across cell types are overall significantly conserved between duplicated genes, compared to random expectations obtained through simulations. However, we show that chromatin contact patterns rapidly diverge with time, to such an extent that gene pairs that originated through duplication more than 300 Mya are indistinguishable from random gene pairs in this respect. In contrast, expression patterns across cell types retain significant similarity for longer periods of time. Finally, we observe a positive correlation between expression pattern divergence and chromatin contact profile divergence, which remains significant when controlling for the association with the age of the duplication event. Our results thus confirm that chromatin contacts play an important role in gene expression regulation.

## Results

### Selection of paralogous genes

We extracted information on paralogous relationships (PRs) between pairs of mouse genes from the Ensembl Compara database, release 102 (Vilella et al. 2009; Harrison et al. 2024). We applied several filters on this PR dataset (Fig. 1A, Materials and methods). First, we kept only PRs where both genes were protein-coding, retaining a total of 354,670 PRs. Second, we discarded PRs where one or both genes were assigned to unplaced contigs or to the mitochondrium, as well as PRs where the two gene coordinates were overlapping. Third, we discarded PRs where one duplicated gene appeared to be derived from a retrotransposition event, reasoning that paralogous genes arisen from such duplication events have very different genomic and regulatory contexts (Carelli et al. 2016), and are thus not suited for a general study of regulatory evolution. Fourth, we implemented a filter to avoid biasing our PR dataset towards large gene families. At this point, 17,664 protein-coding genes are involved in PRs. The number of PRs *per* gene varies between 1 and 373 (Fig. S1), which indicates the presence of large paralogous gene families. To avoid biasing our dataset towards these families, we filtered our dataset keeping only those gene pairs which were each other’s unique most recent paralog. To do this, we analyzed gene tree topologies for each gene family and selected terminal duplication nodes, that is, duplication nodes for which none of their descending nodes corresponded to duplication events in mouse (Fig. 1B, Materials and methods). After this filter, each retained gene appears in exactly 1 PR. Then, we filtered PRs to extract those that could be analyzed without ambiguity with promoter capture Hi-C (PCHi-C) data. For that, we used a PCHi-C dataset collected by Laverré and collaborators (Laverré, Tannier, and Necsulea 2022), which covers 14 samples and 8 cell types for mouse (Table S1). We used RNA-seq data from the same publications, for a similar set of cell types (Table S2).

**Figure 1:**
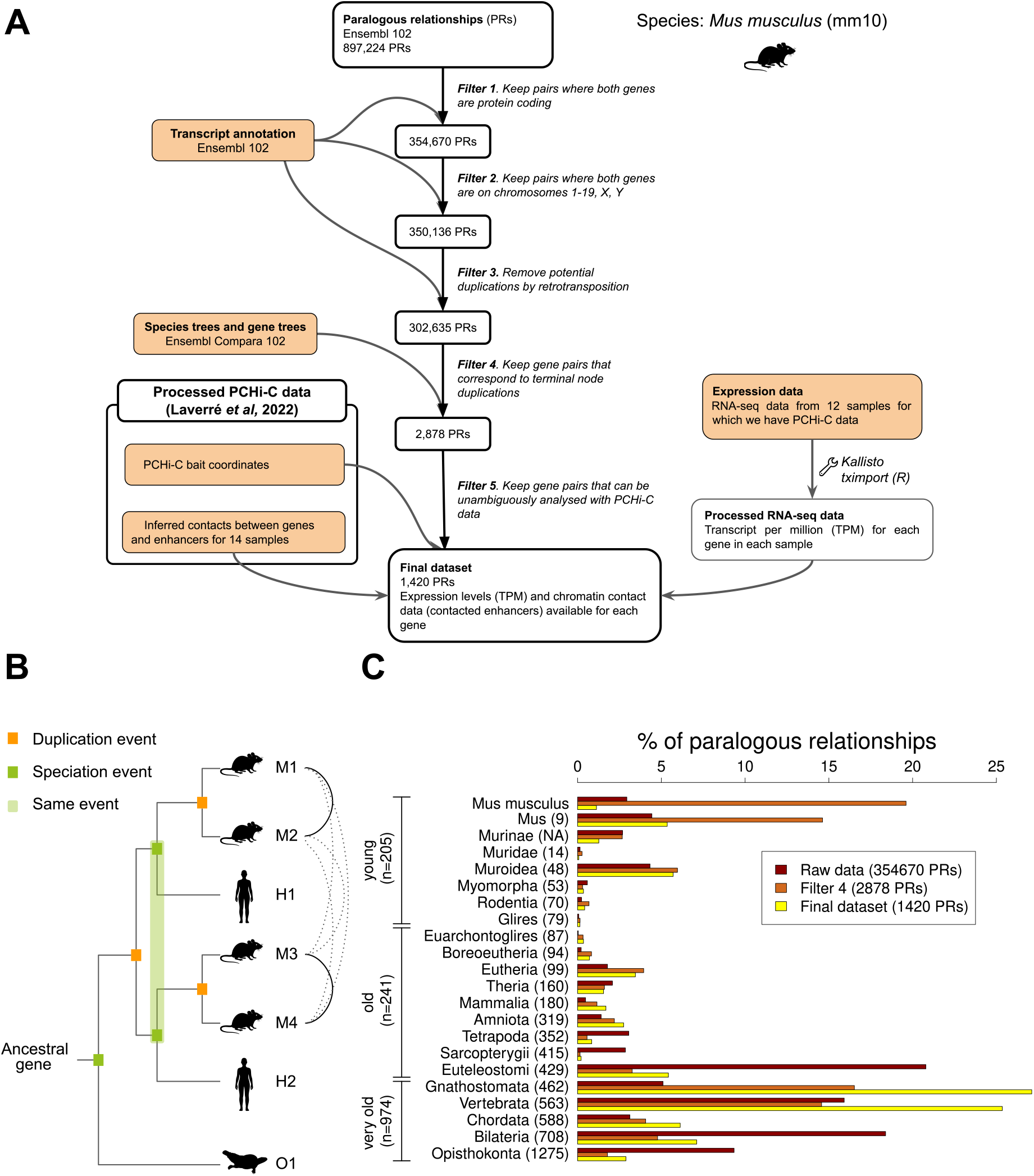
Selection of paralogous relationships and duplication age distribution. **A.** Paralogous gene filtering procedure. **B.** Schematic illustration of the terminal duplication node criterion. Four mouse genes (M1 to M4) are involved in paralogous relationships (PR). For each gene pair, we examine the subtree descending from their last common ancestor: if no additional duplication is present within this subtree, the PR is retained (continuous connecting lines); otherwise, it is discarded (dotted connecting lines). **C.** Histogram representing the distribution of PR duplication ages. The Y-axis indicates the last common ancestor, with divergence times (in million years) retrieved from TimeTree. Several datasets are represented: all PRs involving protein-coding genes (raw data); PRs corresponding to terminal node duplications (filter 4); PRs remaining after the full filtering procedure (final dataset).

As described by Laverré, Tannier, and Necsulea 2022, we filtered PCHi-C chromatin contacts, retaining only those that occurred between genomic regions situated on the same chromosome, at a distance between 25 kilobases (kb) and 2 megabases (Mb). In addition, we retained only those PRs where each paralogous gene promoter was targeted by a single bait and where the two genes were targeted by different baits (Materials and methods). Last, to focus on chromatin contacts between gene promoters and enhancers, we only kept PRs for which each gene contacted at least one ENCODE-predicted enhancer (Materials and methods). Our final dataset consists of 1,420 PRs.

We analyzed the distribution of gene duplication ages, as inferred by Ensembl Compara (Vilella et al. 2009; Harrison et al. 2024), before and after the different filters (Fig. 1C). In the complete set of paralogous relationships within the mouse genome, 75.5% of all duplicated events are estimated to have occurred more than 400 million years ago (Mya; Fig. 1C), and only 14.5% less than 50 Mya. Our fourth filter, which aims to reduce the over-representation of large gene families by selecting only gene pairs which are each other’s unique most recent paralog, enriches the distribution in younger duplication events (Fig. 1C). After this filter, 43.1% of PRs are estimated to have occurred less than 50 Mya and 45.1% more than 400 Mya (Fig. 1C). Conversely, our last filter, which retains gene pairs that are analyzable with PCHi-C data, namely pairs for which both genes have at least one detected interaction in the PCHi-C dataset, shifts the distribution back towards older duplication events: in our final dataset, 13.5% of PRs are estimated to have occurred less than 50 Mya and 74.2% more than 400 Mya (Fig. 1C). Thus, our final set of PRs is similar to the entire set of mouse PRs in terms of duplication age distribution. We note that estimated duplication ages are well correlated with the extent of protein-coding sequence divergence (Fig. S2), as expected.

### Expression and chromatin contact characteristics of paralogous genes

We inferred promoter-enhancer contacts using genome-wide enhancer predictions from ENCODE (Yue et al. 2014) and a PCHi-C dataset generated by various research groups and processed by Laverré, Tannier, and Necsulea 2022. We evaluated gene expression levels using RNA sequencing data that was generated for similar cell types, by the same research groups (Materials and methods).

Given that recent gene duplications have a high degree of sequence similarity, which can give rise to read mapping ambiguity, we first wanted to verify whether we could adequately estimate the expression levels of the protein-coding genes included in our PR dataset. To do this, we simulated single-end and paired-end RNA-seq reads, using a realistic gene expression level distribution (Materials and methods). We then estimated gene expression levels with kallisto (Bray et al. 2016) and compared the estimated levels with the true simulated values (Fig. S3). We show that gene expression levels predicted with kallisto are highly correlated with the true simulated values across all protein-coding genes (Spearman’s *ρ* = 0.99 for single-end data and 0.96 for paired-end data, Fig. S3). Importantly, we observed similar expression level correlation values for the 2,840 paralogous genes included in our final dataset (Spearman’s rho = 0.99 for single-end data and 0.97 for paired-end data, Fig. S3). Moreover, we observed that the difference in expression levels between the two genes in a PR is well correlated between the simulated values and the estimates obtained with kallisto (Spearman’s *ρ* = 0.96 for single-end data and 0.9 for paired-end data across the 1,420 PRs, Fig. S4). Importantly, the correlation values were similar for young PRs (estimated duplication age *≤* 79 Mya, Fig. S3, Fig. S4). These results validate our capacity to evaluate gene expression for duplicated genes, and in particular for young PRs.

We note that duplicated gene characteristics vary with the age of duplication. In particular, the average expression level and the number of contacts increase with the age of duplication (Fig. S5). Likewise, the genomic localization of gene pairs varies with the duplication age: young duplicated gene pairs (estimated duplication age *≤* 79 Mya) are localized in *cis* in 88% of the cases, while old paralogs (duplication age superior to 79 Mya and inferior to 462 Mya) are more evenly distributed, with 53% of gene pairs in *cis*, and very old paralogs (duplication age *≥* 462 Mya) are predominantly localized in *trans*, in 91% of the cases (Fig. S5, S6). This observation is consistent with the presence of old paralogs derived from whole genome duplications in the vertebrate ancestor (Dehal and Boore 2005). Earlier duplications are likewise overwhelmingly localized in *trans* (Fig. S6), as expected if, with time, segmental duplications are broken apart by subsequent genomic rearrangements (Lan and Pritchard 2016).

Finally, we note that gene biological functions also vary with the age of duplication. Gene Ontology analyses revealed an enrichment for biological processes related to cellular metabolism (including RNA metabolism, noncoding RNA processing, translation *etc.*) for older duplicated genes, and an enrichment for sensory processes (including perception of smell) and immune response for younger duplicated genes (Tables S3 and S4, Materials and methods). This pattern is consistent with the presence of olfactory receptor genes among the younger duplicates in mouse (Niimura and Nei 2006), and with the presence of highly expressed genes among the older duplicates, as discussed above.

### Construction of simulated datasets

To test if paralogous gene characteristics are different from random expectations, we constructed two types of simulated control datasets. The first control (hereafter called shuffled dataset) was generated by shuffling paralogous genes, creating random gene pairs. The second control (hereafter called sampled dataset) was generated by randomly sampling gene pairs among genes that are not part of PRs, in such a way as to obtain similar distributions compared to real PRs in terms of average expression levels, of relative genomic localization (*cis* or *trans*), and (for *cis* pairs) of distances between genes (Materials and methods). We confirmed *a posteriori* that the sampled datasets are overall similar to the real PRs in terms of average expression level distribution (Fig. S7). We drew 1000 simulated datasets of each type.

In the analyses presented below, we compare the distributions observed for the real PRs with the distributions observed for a typical shuffled or sampled dataset (Materials and methods), for various expression divergence and chromatin contact divergence metrics. We also compare median divergence values between the PRs and the 1000 simulated datasets.

### Evolutionary divergence of expression levels and regulatory complexity

We first aimed to evaluate if average gene expression levels and total numbers of contacted enhancers are conserved between paralogous genes. To do this, we used two simple metrics: first, we defined Δexpression as the difference in expression levels (averaged across all biological samples) between the two genes in a PR, normalized by dividing by the average expression level of the two genes. Second, we defined Δcomplexity as the difference in the numbers of contacted enhancers (considered across all biological samples) between the two genes in each PR, divided by the average number of contacted enhancers for the two genes (Materials and methods). This latter metrics is named Δcomplexity because the number of enhancers associated with a given gene can be seen as a measure of the gene’s regulatory complexity (Berthelot et al. 2018). We note that Δexpression is negatively correlated with the minimum expression level and positively correlated with the maximum expression level of the two genes in the PR (Fig. S8), as expected given that it is computed from the difference between the two. Likewise, Δcomplexity is significantly correlated with the minimum and maximum number of contacted enhancers of the two genes in the paralogous pair (Fig. S9). In contrast, Δexpression is not significantly correlated with the average expression level and Δcomplexity is not correlated with the mean number of enhancers contacted by the two genes in the paralogous pair (Fig. S8 and Fig. S9).

We first compared Δexpression and Δcomplexity between PRs and simulated datasets. We observed that Δexpression is much smaller for PRs than for control datasets (Wilcoxon test for the comparison with a typical control dataset, *p*-value 9.8 *×* 10*^−^*^28^ for the comparison with the sampled dataset, 2.4 *×* 10*^−^*^46^ for the comparison with the shuffled dataset, Fig. 2), while Δcomplexity is only slightly smaller for PRs than for shuffled and sampled datasets (Wilcoxon test for the comparison with a typical control dataset, *p*-value 4.7 *×* 10*^−^*^7^ and 0.034, respectively, Fig 2). Consistently, the median Δexpression value is lower for PRs than the median values observed for all 1000 shuffled and sampled control datasets, while the median Δcomplexity for PRs is higher than the median values observed for 22 out of 1000 sampled datasets (Fig. S10). These observations suggest that average expression levels are highly conserved between duplicated genes, more so than numbers of contacted enhancers.

**Figure 2:**
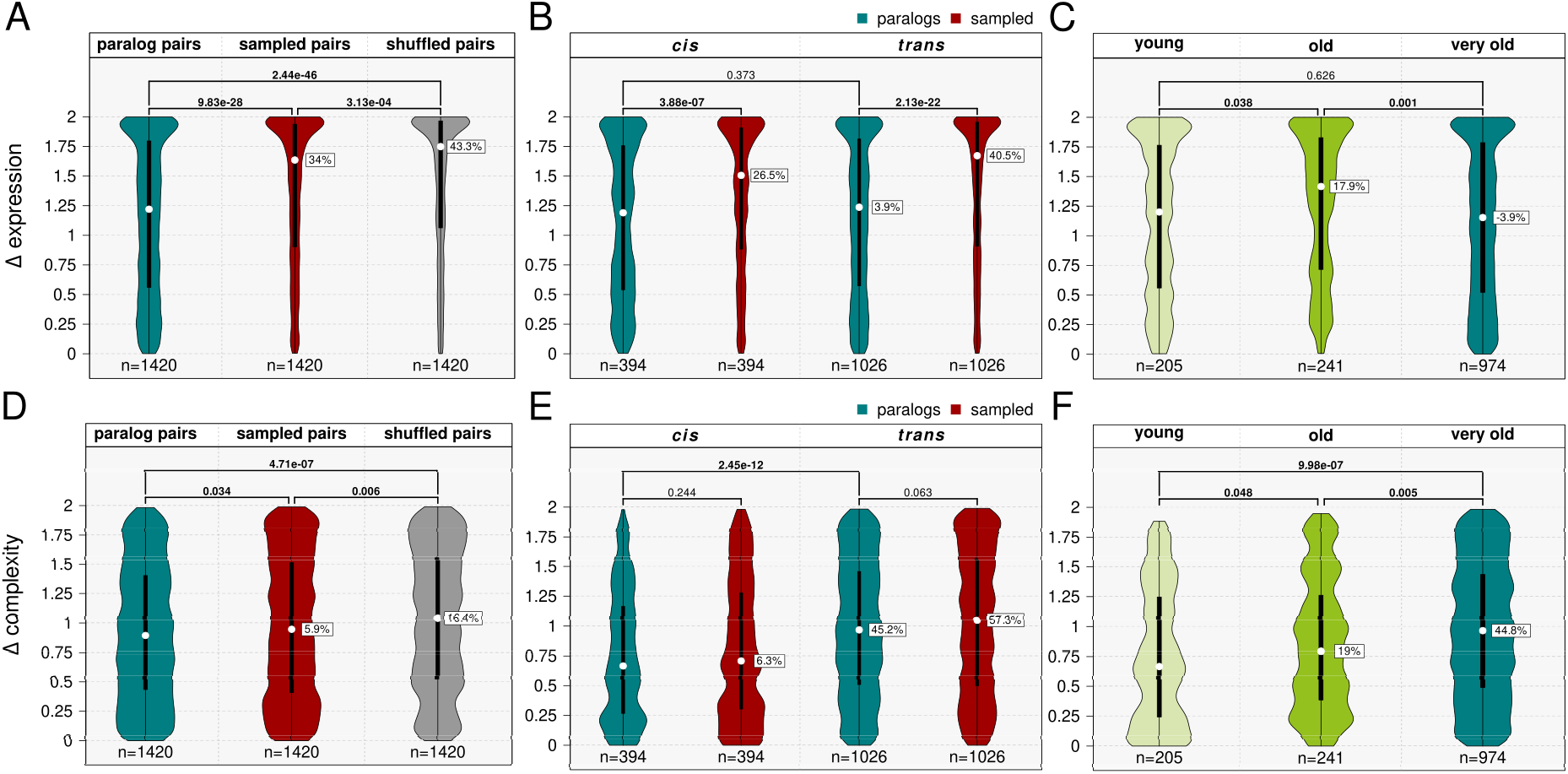
Evolutionary conservation of expression levels and regulatory complexity. **A.** Blue-green: violin plot showing the distribution of the normalized difference in mean expression levels (Δexpression) between paralogous genes. Red: same, for randomly sampled gene pairs, selected to match the paralogs in both expression-level distribution and distance between gene transcription start sites (TSS), as well as chromosomal localization (Materials and methods). Gray: same, for shuffled paralogous gene pairs. **B.** Same as **A**, with datasets separated according to gene localization (*cis*/*trans*), for real and randomly sampled paralogous genes. **C.** Violin plot showing Δexpression with paralogs pairs separated according to the approximate age of their duplication event: *young* corresponds to duplications that occurred from *Mus musculus* to *Glires*, *old* from *Euarchontoglires* to *Euteleostomi*, and *very old* to duplications older than *Gnathostomata*. **D–F.** Same as **A–C**, for the normalized difference in the total number of enhancers contacted by each gene (Δcomplexity). For all panels, *p*-values correspond to Wilcoxon tests. *P*-values lower than 0.05 are indicated in bold. Percentages indicated in white rectangles correspond to relative differences between the medians of the leftmost violin plot and the labelled violin plot. **A-B.** The random seeds used to generate the typical sampled and shuffled control datasets displayed in the figure were 367 and 223, respectively. **D-E.** The random seeds used to generate the typical sampled and shuffled control datasets displayed in the figure were 759 and 824, respectively.

We next asked whether Δexpression and Δcomplexity are associated with gene co-localization. There was no significant difference in Δexpression between *cis*-localized and *trans*-localized gene pairs (Wilcoxon test, *p*-value 0.374, Fig. 2), but *cis*-localized pairs had much lower Δcomplexity values than *trans*-localized pairs (Wilcoxon test, *p*-value 2.45 *×* 10*^−^*^12^, relative median difference 45.2%, Fig. 2). Moreover, gene pairs separated by at most 1 Mb on the same chromosome had significantly lower Δcomplexity values than more distant *cis*-localized pairs (Wilcoxon test, *p*-value 0.018, Fig. S11), while no significant difference between the two categories was observed for Δexpression. We note that *trans*-localized pairs include genes derived from whole-genome duplication events, also called *ohnologs* in reference to Susumu Ohno (Ohno 1970). We found no difference between *ohnologs* and non-*ohnologs* among *trans*-localized PRs, neither for Δexpression nor for Δcomplexity (Fig. S12). Our results show that duplicated genes found on the same chromosome and in particular those found in close proximity to each other have similar numbers of contacted enhancers. This is due at least in part to the fact that they contact common enhancers: compared to control datasets, genes in real PRs have larger proportions of common contacted enhancers (Fig. S13, Materials and methods), as previously observed (Ibn-Salem, Muro, and Andrade-Navarro 2017).

We then tested if there was an association between duplication age, on one hand, and Δexpression and Δcomplexity, on the other. To do this, we divided PRs into three age classes (Fig. 1): “young” PRs, which originated in the Glires ancestor or later (estimated duplication age *≤* 79 Mya); “old” PRs, which originated in ancestors of groups ranging from Eurarchontoglires to Euteleostomi (estimated duplication age 87 to 429 Mya); and “very old” duplications, which originated in the Gnathostomata ancestor or earlier (estimated duplication age 462 to 1275 Mya). We found that the age of duplication is not linearly correlated with Δexpression (Fig. 2C). Old PRs had slightly but significantly higher Δexpression values than the two other age classes (Fig. 2C, Wilcoxon test *p*-value = 0.038 and 0.001 for old *vs* young: and old *vs* very old, respectively). These differences become weaker when separating duplicated genes by both duplication age and genomic co-localization (Fig. S14). Furthermore, we observe that duplication age is associated with Δcomplexity, with an effect size of around 20% between successive age groups (Fig. 2F, *p*-value 0.048 for the comparison between “young” and “old” PRs, *p*-value 0.005 for the comparison between “old” and “very old” PRs). This association is consistent with expectations, as older PRs have higher Δcomplexity than younger PRs. However, for *cis* and *trans* pairs there was no significant difference between PR age classes (Fig. S14). Given that the genomic localization is strongly associated with the duplication age, the relationship between Δcomplexity and duplication age needs to be interpreted with caution, as genomic localization may be a confounding factor.

Finally, we investigated the relationship between Δexpression and Δcomplexity, across PRs. It was previously shown that gene expression levels are positively correlated with regulatory complexity, defined as the number of enhancers associated with each gene (Berthelot et al. 2018; Laverré, Tannier, and Necsulea 2022). We confirm this observation for all protein-coding genes as well as for genes involved in PRs (Fig. S15). From this correlation, it follows that differences in expression levels should be positively correlated with differences in numbers of contacted enhancers, across gene pairs - whether they are paralogous or not. Indeed, we observed a weak positive correlation between Δexpression and Δcomplexity across PRs (Fig. 3A–B, Spearman’s *ρ* = 0.07, correlation test *p*-value 0.011). However, this correlation was significantly weaker than those obtained for the 1000 sampled and shuffled datasets: for the shuffled datasets, only 24 out of 1000 simulations showed lower *ρ* values than the PRs, while for the sampled datasets, only 8 simulations showed lower *ρ* values (Fig. 3C). We hypothesize that this may be due to a strong constraint on average expression levels (consistent with our observations for Δexpression), and to a weaker constraint on the numbers of contacted enhancers (consistent with our observations for Δcomplexity). This may result in an uncoupling of the rates of expression level evolution and of regulatory complexity evolution, and thus in a weaker than expected correlation between Δexpression and Δcomplexity.

**Figure 3:**
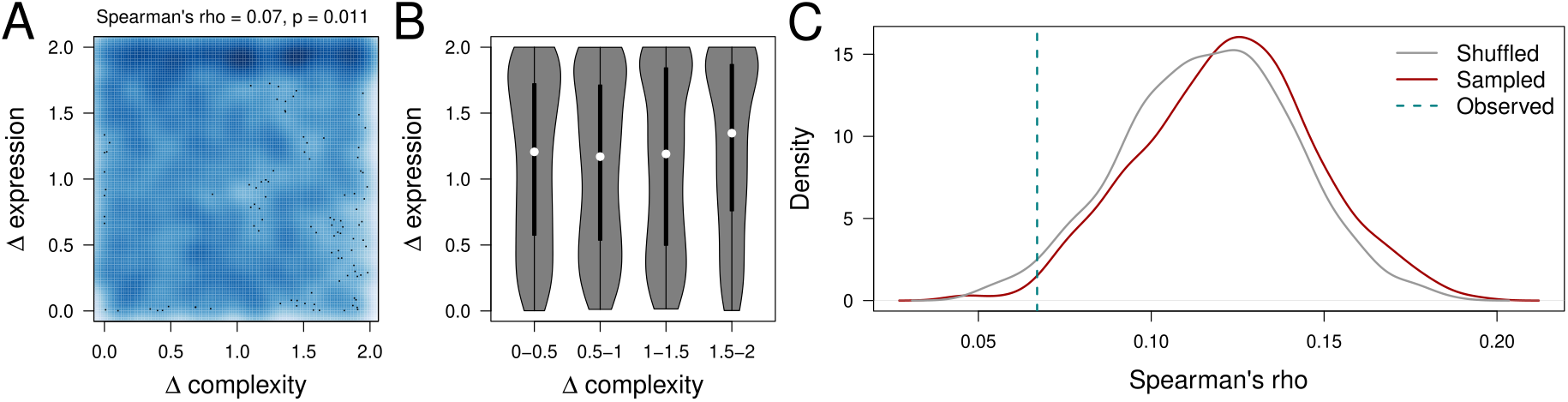
Relationship between Δexpression and Δcomplexity. **A.** Smooth scatter plot representing the relationship between Δcomplexity (X-axis) and Δexpression (Y-axis). **B.** Violin plots showing the distribution of Δexpression values, across 4 classes of PRs defined based on Δcomplexity (0 to 0.5, 0.5 to 1, 1 to 1.5 and 1.5 to 2). **C.** Density plot showing the distribution of Spearman’s correlation coefficients (*ρ*) between Δexpression and Δcomplexity values, from 1000 sampled and 1000 shuffled datasets. The dotted vertical line represents the value observed for the PR dataset. Only 8 of the sampled datasets and 24 of the shuffled datasets show Spearman’s *ρ* values lower than the one observed for PRs.

### Evolutionary divergence of expression profiles and chromatin contact profiles of duplicated genes

The two metrics described above (Δexpression and Δcomplexity) were designed to evaluate the evolution of the average gene expression levels and of the total number of enhancers contacted by genes, combined across all available biological samples. To gain a more detailed insight into the evolution of expression and regulatory contact patterns, we designed additional measures of expression and regulatory divergence that take into account variations across biological samples. We first defined relative expression profiles, by dividing gene expression levels (measured as TPM) by the maximum value observed across all samples, for each gene (Materials and methods). Then, for each pair of paralogous genes, we defined the expression profile divergence as the Manhattan distance between the relative expression profiles of the two genes (Materials and methods). To measure chromatin contact profile divergence, we first counted for each gene the number of ENCODE-predicted enhancers that it contacts in each biological sample. We then transformed these counts into frequencies (*per* million) across all genes for each sample, obtaining a metric called EPM (enhancers *per* million). As for the TPM, we then computed relative EPM values by dividing the EPM values by the maximum EPM value observed across all biological samples, separately for each gene. We defined the contact profile divergence as the Manhattan distance between the relative EPM profiles for the two genes in a PR (Materials and methods). We note that this measure of contact profile divergence does not test whether the same enhancers are contacted in different cell types.

Consistent with our observations for Δexpression, we found that the expression profile divergence was significantly lower for PRs than for shuffled and sampled datasets (Fig. 4A, Wilcoxon test, *p*-value 1.41 *×* 10*^−^*^31^ for the comparison between PRs and a typical sampled dataset, relative difference 20%). The expression divergence was lower for PRs than for all 1000 sampled and shuffled datasets (Fig. S16). The same was true for the contact profile divergence, although the difference between PRs and simulated datasets was lower than for the expression profile divergence (Fig. 4D, Wilcoxon test, *p*-value 1.08 *×* 10*^−^*^4^ for the comparison between PRs and a typical sampled dataset, relative difference 4.3%).

**Figure 4:**
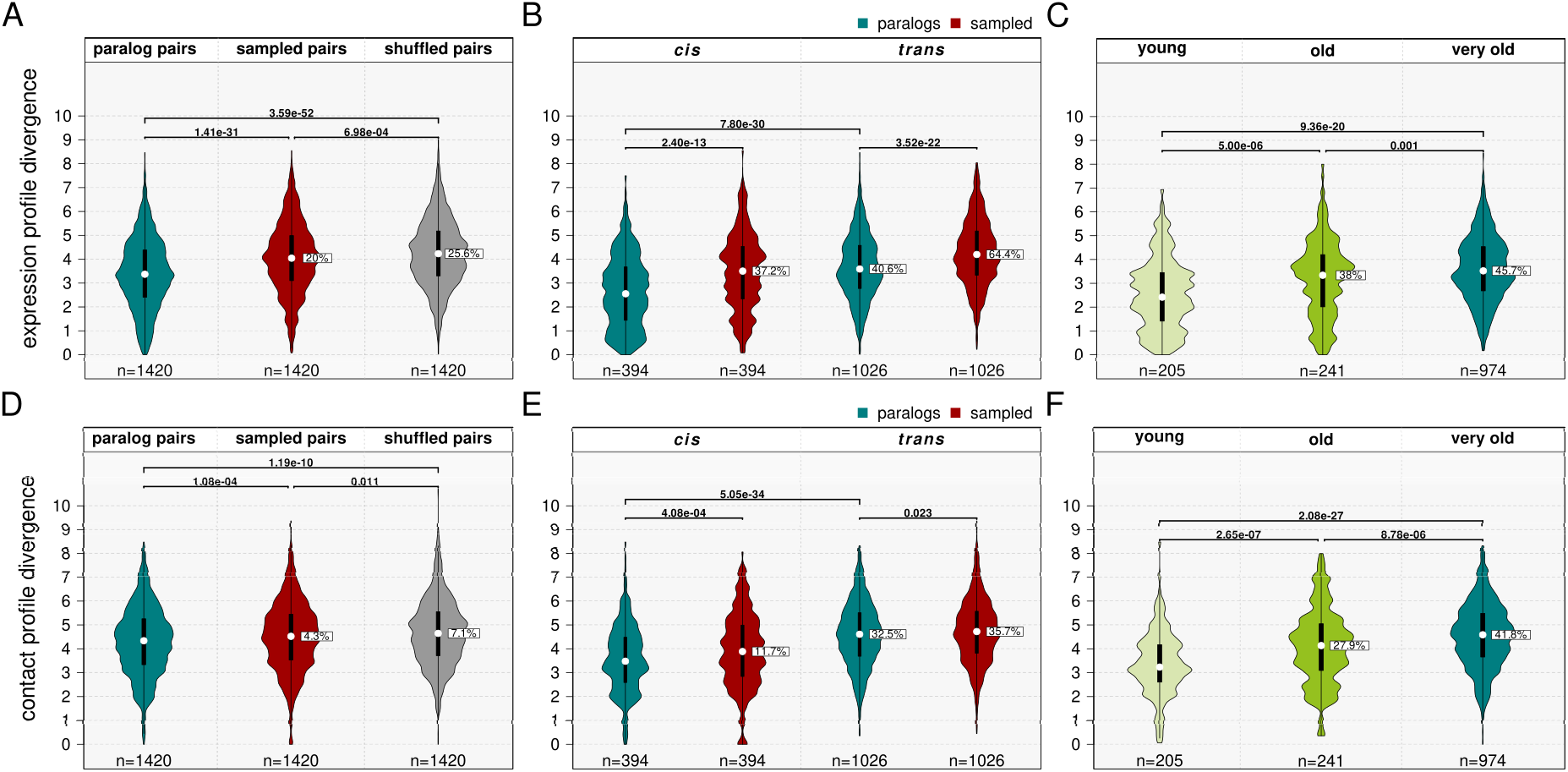
Conservation of expression profiles and of chromatin contact profile. **A.** Violin plot showing the distribution of the expression profile divergence for pairs of paralogous genes (blue-green) and control datasets. The sampled dataset (red) corresponds to gene pairs selected to match the paralogs for average expression levels, distance between transcription start sites (TSS) and chromosomal localization. The shuffled dataset (gray) corresponds to a permutation of the paralog pairs. **B.** Same as **A**, but with datasets separated according to gene localization (*cis*/*trans*). **C.** Violin plot showing the divergence of the expression profile with paralogs pairs separated according to the approximate age of their duplication event: *young* corresponds to duplications that occurred from *Mus musculus* to *Glires*, *old* from *Euarchontoglires* to *Teleostomi*, and *very old* to duplications older than *Gnathostomata*. **D–F.** Same as **A–C**, for the divergence of chromatin contact profiles. For all panels, *p*-values correspond to Wilcoxon tests and *p*-values represented in bold are below 0.05. Percentages indicated in white rectangles correspond to the relative difference between the medians of the leftmost violin plot and the labelled violin plot. **A-B.** The random seeds used to generate the typical sampled and shuffled control datasets displayed in the figure were 727 and 174, respectively. **D-E.** The random seeds used to generate the typical sampled and shuffled control datasets displayed in the figure were 48 and 261, respectively.

In contrast with what we observed for Δexpression and Δcomplexity, we found that the genomic localization is significantly associated with both the expression divergence and the contact profile divergence. Specifically, we observed lower divergence values for *cis*-localized pairs than for *trans*-localized pairs (Fig. 4B,E), and for *cis*-localized genes found in close proximity in the genome (distance *≤* 1Mb) than for more distant *cis*-localized gene pairs (Fig. S17). Furthermore, for *trans*-localized genes, we observed significant lower expression profile divergence values for non-*ohnologs* than for *ohnologs* (Fig. S18). For the contact profile divergence, there was no difference between these two PR categories.

Finally, we examined the relationship between the age of duplication, on one hand, and the expression profile divergence and the contact profile divergence, on the other. We found a significant positive association between the expression profile divergence and the age of duplication (Fig. 4C), as well as between the contact profile divergence and the age of duplication (Fig. 4F). The tendency remained the same when analyzing *cis* and *trans* PRs separately, although the differences between consecutive age classes were not always statistically significant (Fig. S19).

### Relationship between the expression profile divergence and the chromatin contact profile divergence

Next, we investigated the association between expression profile divergence and regulatory profile divergence. We observed a weak but significant correlation between the two variables across PRs (Fig. 5A–B; Spearman’s *ρ* = 0.17, *p*-value 1.12 *×* 10*^−^*^10^). The correlation observed among PRs was overall higher than the correlations obtained from the 1000 sampled and shuffled datasets. Specifically, none of the shuffled datasets exhibited a higher *ρ* value, and only 72 of the 1000 sampled datasets showed a higher value than the one observed for PRs (Fig. 5C). This observation is in contrast to our findings on the relationship between Δexpression and Δcomplexity.

**Figure 5:**
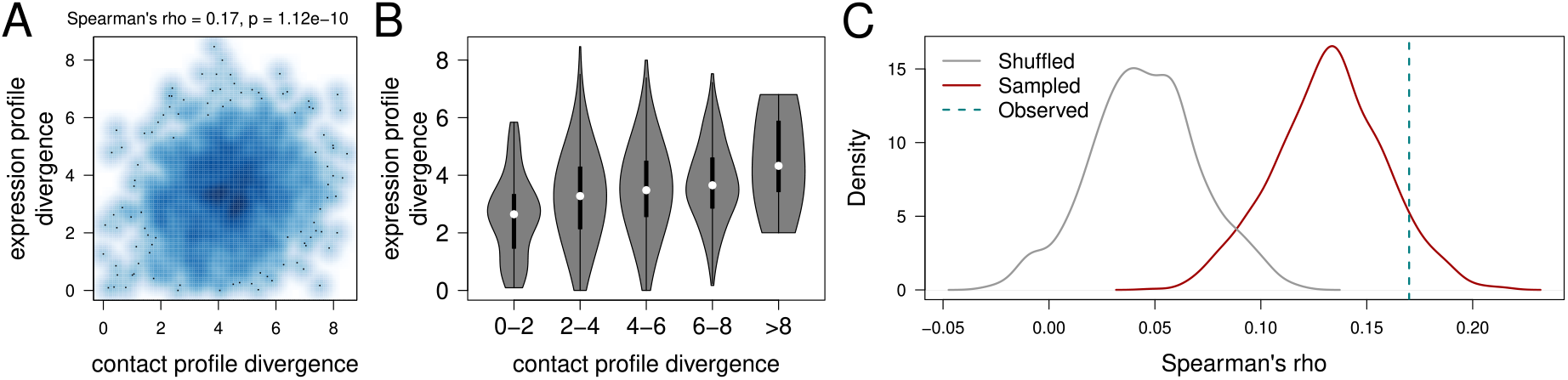
Relationship between the expression profile divergence and the chromatin contact profile divergence. **A.** Smooth scatter plot showing the relationship between the expression profile divergence and the contact profile divergence, across paralogous gene pairs. **B.** Violin plot showing the distribution of the expression profile divergence, across classes of genes defined based on the contact profile divergence. **C.** Density plot showing the distribution of Spearman’s correlation coefficient (*ρ*) between expression profile divergence and contact profile divergence, derived from 1000 sampled and shuffled datasets. Only 72 out of 1000 sampled datasets and none of the shuffled datasets show a *ρ* value higher than that of the real PR dataset.

The positive correlation between the expression profile divergence and the chromatin contact profile divergence, across PRs, is consistent with a causal relationship between changes in chromatin contact profiles and gene expression evolution. Indeed, in the simplest evolutionary scenario, mutations that induce regulatory landscape changes (including chromatin contact alterations) are likely to trigger expression profile changes. However, we note that the two variables are correlated with the time since the duplication event, as well as with other factors, as discussed above. Notably, both expression profile divergence and contact profile divergence are correlated with the rate of non-synonymous substitutions in protein-coding sequences (Spearman’s *ρ* 0.13 in both cases, correlation test *p*-value *<* 1.5 *×* 10*^−^*^6^, Fig. S20), which is a good proxy for the time since the duplication event (Fig. S2). Moreover, we find a significant positive correlation between the expression profile divergence and the mean or maximum expression level of the two genes involved in a PR (Spearman’s *ρ* 0.16 for the mean expression level, Spearman’s *ρ* 0.18 for the maximum expression level, correlation test *p*-value *<* 2.2 *×* 10*^−^*^9^, Fig. S21). Significant positive correlations are likewise observed between the chromatin contact profile divergence and the minimum, mean or maximum number of contacted enhancers for the two genes involved in a PR (Spearman’s *ρ* 0.11, 0.31 and 0.33, for the minimum, mean and maximum, respectively; correlation test *p*-value *<* 2.7 *×* 10*^−^*^11^, Fig. S22). Furthermore, we also observed that the genomic localization is significantly correlated with our expression and contact divergence measures (see above). To test whether the correlation between the expression profile divergence and the contact profile divergence remains positive and significant when these factors are taken into account, we constructed a linear regression that models the relationship between the expression divergence on one hand, and the maximum expression level, the non-synonymous substitution rate, the genomic localization and the chromatin contact profile divergence, on the other hand (Table 1). We did not include the number of contacted enhancers as an explanatory variable because it is not independent from the gene expression level, as shown above (Fig. S15). We found that the linear regression coefficients were significantly different from 0 for all considered explanatory variables. In particular, the association between expression profile divergence and contact profile divergence remains significant and positive (Student’s test, *p*-value 0.0045, Table 1).

**Table 1:**
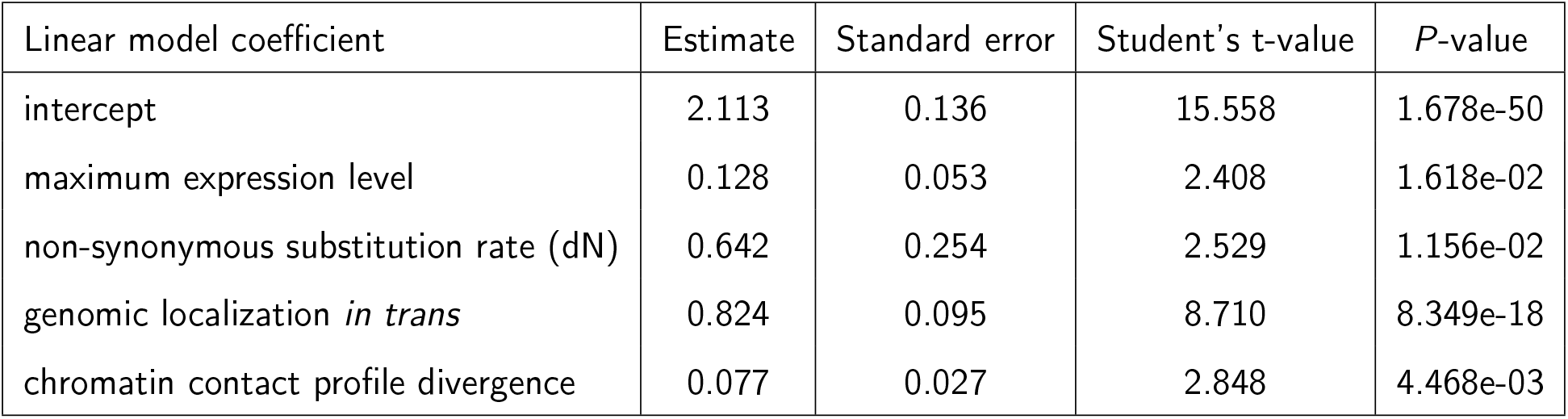
Results of a linear regression that models the relationship between the expression profile divergence, on one hand, and the maximum gene expression level (expressed as log-transformed TPM), the non-synonymous substitution rate, the chromatin contact profile divergence and the genomic localization (*cis* / *trans*), on the other hand. The regression is performed across the 1,420 PRs.

### Temporal dynamics of the expression profile and contact profile divergence

Our last aim was to understand how expression divergence and chromatin contact divergence accumulate with time. We examined the temporal dynamics of the expression profile divergence and the regulatory profile divergence, by comparing paralogous gene pairs with different duplication ages (Fig. 6). Paralogous gene pairs span a broad range of estimated divergence times, between *≈*9 Mya (for gene pairs whose last common ancestor was in the *Mus* genus) and *≈*1275 Mya (for gene pairs whose last common ancestor corresponds to the ancestor of the *Opisthokonta* clade). We grouped PRs by divergence times and analyzed groups represented by at least 20 gene pairs. Both the expression profile divergence and the contact profile divergence have a non-linear association with the divergence time. Namely, for divergence times below 300 My, recently diverged gene pairs have lower divergence values than older gene pairs, as expected (Fig. 6). Within this interval, both measures show a departure from linearity with a high divergence value observed for gene pairs that originated in the Therian ancestor (estimated divergence time 160 Mya) (Fig. 6). We note that this age group and the following one (genes originating in the Mammalian ancestor, estimated divergence time 180 Mya) have the smallest sizes, with only 22 and 24 gene pairs, respectively. Beyond 300 My, the expression profile divergence is negatively associated with divergence time (Fig. 6A), while the contact profile divergence reaches a plateau (Fig. 6B). Moreover, the expression profile divergence is smaller for paralogous gene pairs than for simulated pairs at all divergence times (Fig. 6A). In contrast, the contact profile divergence is similar to the values observed for simulated datasets, for gene pairs having originated 300 Mya or earlier (Fig. 6B).

**Figure 6:**
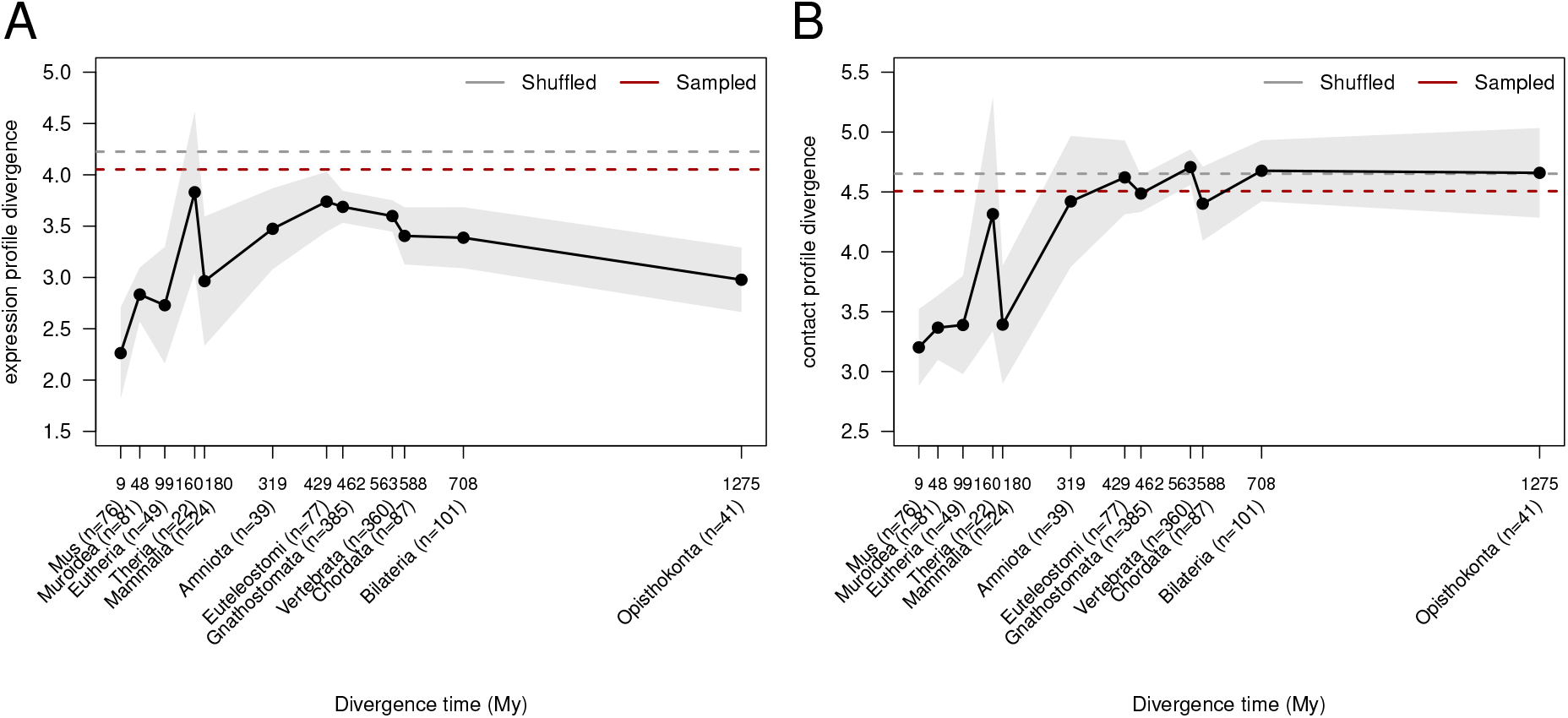
Dynamics of the expression divergence and contact divergence with time. **A.** Distribution of the median expression profile divergence as a function of the estimated divergence time of paralogous gene pairs. Sampled median value: 4.05. Shuffled median value: 4.22 **B.** Distribution of the median contact profile divergence as a function of the estimated divergence time of paralogous gene pairs. Sampled median value: 4.51. Shuffled median value: 4.65 **A,B.** Dots represent median values. Gray shading represents 95% confidence intervals of the median. Dotted lines represent the median values observed for shuffled datasets (gray) and sampled datasets (red). X-axis labels represent the last common ancestor inferred for PRs and the corresponding estimated divergence times, in My. The numbers of gene pairs included in each group are given in parentheses.

We observed similar temporal dynamics patterns for our other measures of expression and contact divergence, namely Δexpression and Δcomplexity (Fig. S23). Namely, Δexpression increases rapidly with time for gene pairs having diverged less than 180 Mya, then decreases with time. The highest Δexpression values are observed for divergence times between 160 and 319 Mya, where they are similar to random expectations derived from sampled and shuffled datasets (Fig. S23A). The temporal dynamics is more irregular for Δcomplexity, which seems to reach a plateau after 319 Mya, similar to the contact profile divergence (Fig S23).

We note that the divergence of protein sequences and of promoter sequences does not follow the same pattern (Fig. S24). The median rate of non-synonymous substitutions between paralogous genes tends to increase with divergence time, with a rapid increase observed for the youngest divergence times and a slower increase for the oldest time points (Fig. S24). For divergence times below 100 Mya, for which homologous promoters could be unambiguously identified for paralogous gene pairs (Materials and methods), we observed a linear increase in promoter sequence divergence with time (Fig. S24). Given that promoter sequences can be aligned between paralogous genes in only 556 gene pairs, we defined a different sequence similarity measure, based on the presence of shared transcription factor (TF) binding motifs in the two promoter sequences (Materials and methods). We observed that duplicated genes shared TF binding motifs more often than randomly sampled gene pairs (Fig. S25), as previously shown (Fraimovitch and Hagai 2023). This measure of promoter sequence similarity has a non-linear association with duplication time (Fig. S25). Thus, promoter sequences do not appear to evolve at similar rates as promoter-enhancer contacts, for duplicated genes.

## Discussion

In this manuscript, we address the evolution of regulatory chromatin contacts that take place between gene promoters and enhancers, for duplicated genes. This aspect of gene expression regulation remains largely unexplored from an evolutionary standpoint, likely due to the absence of comparable, high-resolution chromatin contact data for different species. Here, our main objective was to obtain a broad overview of regulatory chromatin contact evolution following gene duplication.

We employed several metrics to study gene expression divergence and chromatin contact divergence between pairs of paralogous genes, and we generated simulated paralogous gene datasets to obtain random expectations. We obtain several encouraging results for the validity of our analyses. First, our Δexpression analyses show that average expression levels are more conserved between paralogous genes than between randomly paired genes. Likewise, our

Δcomplexity analyses show that the total numbers of contacted enhancers are more similar for paralogous genes than for randomly paired genes. Similar results were obtained for our measure of the divergence in relative expression levels across cell types, and for our measure of the divergence in relative numbers of contacted enhancers across cell types. Thus, we are confident the metrics that we employed are able to capture biologically relevant information pertaining to gene expression and regulatory evolution.

Our results indicate that chromatin contacts between promoters and enhancers evolve considerably faster than gene expression, across duplicated genes. Indeed, although we cannot directly compare the expression divergence and contact divergence metrics, we observe stronger contrasts between paralogous genes and randomly paired genes for gene expression than for chromatin contacts. Moreover, expression divergence and chromatin contact divergence display different temporal dynamics, with the latter reaching a plateau that coincides with the random expectation after approximately 300 My, while the former remains below random expectations for all divergence times. These results are consistent with previous observations obtained through comparisons across species, for other aspects of gene expression regulation such as enhancer sequences and activity or transcription factor binding (Wong et al. 2015; Berthelot et al. 2018), as well as for promoter-enhancer chromatin contacts (Laverré, Tannier, and Necsulea 2022). This discrepancy between regulatory evolution and expression evolution may be explained by the robustness of gene expression regulation, which involves numerous *cis*-regulatory elements and *trans*-acting factors. Individual regulatory changes are thus not expected to have dramatic consequences on gene expression, suggesting that rapid regulatory evolution can occur while gene expression remains well conserved.

Importantly, we found that, across duplicated gene pairs, the divergence of expression profiles across cell types is significantly and positively correlated with the divergence of chromatin contact profiles. This result is notable because correlations between regulatory evolution and expression evolution have long been elusive (Wong et al. 2015; Berthelot et al. 2018). We note that a weak but significant positive correlation between expression divergence and chromatin contact divergence, across genes, was previously reported for a comparison between human and mouse (Laverré, Tannier, and Necsulea 2022). Thus, genes that have high rates of evolution in terms of chromatin contacts also have high rates of expression evolution, which is consistent with a significant effect of chromatin contact changes on gene expression. Here, we confirm this previous report, but with a *caveat*: our correlations between expression divergence and chromatin contact divergence are measured across duplicated genes, which have highly variable divergence times. Thus, if both expression divergence and chromatin contact divergence are positively correlated with the age of the duplication event, we expect a positive correlation between the two even in the absence of a causal relationship between regulatory changes and expression changes. To account for this possible confounding factor, we performed multivariate analyses that test for an association between expression divergence, on one hand, and chromatin contact divergence, protein sequence divergence (used as a proxy for the time since the duplication event) as well as other potential confounding factors such as genomic localization or expression levels, on the other hand. Reassuringly, the association between expression divergence and chromatin contact divergence remains significant even when these additional confounding factors are explicitly included in the model. This observation reinforces the association between expression divergence and chromatin contact divergence previously reported for orthologous genes between human and mouse (Laverré, Tannier, and Necsulea 2022), suggesting that chromatin contacts play an important role in gene expression regulation.

We believe that many of the results that we obtained for duplicated genes reflect general patterns in the evolution of chromatin contacts and into its link with expression evolution. Nevertheless, our study also revealed several patterns that are undeniably unique to duplicated genes. First, we observe a strong association between chromatin contact divergence and genomic localization of paralogous genes. Specifically, we find that duplicated genes that are localized in *cis* (*i.e.* on the same chromosome) are more conserved than duplicated genes that are localized in *trans*, in terms of the total number of contacted enhancers and of chromatin contact profiles across samples. Relative expression profiles across cell types are likewise more conserved for *cis* gene pairs than for *trans*, although we do not see this pattern for Δexpression. Moreover, we observed that paralogous genes found close together on the same chromosome are more similar than distant gene pairs, in terms of average numbers of contacted enhancers, of expression profiles and chromatin contact profiles across samples. For gene expression profiles, our results confirm previous reports that expression sub-functionalization and neo-functionalization are more frequent for genes localized on different chromosomes (Lan and Pritchard 2016). It was previously proposed that chromosomal separation of duplicated genes is necessary to achieve regulatory divergence and that tandem duplicates tend to be co-regulated (Lan and Pritchard 2016). Our results bring evidence that this is indeed the case, and that promoter-enhancer chromatin contacts are part of the conserved regulatory mechanisms of *cis*-localized duplicates.

Our observation that expression profiles and chromatin contact profiles have different temporal dynamics, through evolutionary time, may likely also be explained by the unique characteristics of duplicated genes. The non-linear pattern of gene expression divergence, with a maximum divergence value observed for duplication events that originated around 300 Mya, was previously observed for duplicated genes (Guschanski, Warnefors, and Kaessmann 2017). A likely explanation for the low expression divergence observed for ancient duplicated genes lies in these genes’ biological characteristics. Indeed, we observed that these genes are enriched in functions related to cellular metabolism, RNA processing, translation *etc*, which is consistent with their high expression levels. It was previously shown that highly expressed genes are preferentially retained in two copies following whole genome duplication events (Gout et al. 2010). For these genes, expression level changes is likely to be costly (Gout et al. 2010), which can explain their low expression divergence values.

Our work paves the way for additional research directions, that would further our understanding of the evolution of regulatory chromatin contacts for duplicated genes. In particular, we note that we were unable to investigate the evolutionary origin of the enhancers that are in chromatin contact with duplicated genes. Indeed, our analyses show that duplicated gene promoters cannot be well aligned for paralogous relationships that originated more than 160 Mya. Enhancer sequences show levels of conservation similar to those of promoter sequences (Villar et al. 2015), although we note that this may depend on the tissues and developmental stages in which they are detected (Blow et al. 2010). Identifying paralogous enhancers is thus likely to be impossible for genes that diverged more than 160 Mya, which constitute a large fraction of the paralogous relationships that we analyzed here. We also did not address the regulatory relationships that may take place between the promoters of *cis*-duplicated genes. Analyses of genome-wide Hi-C data previously revealed that paralogous genes are involved in long-distance chromatin contacts more often than expected by chance (Ibn-Salem, Muro, and Andrade-Navarro 2017). In this study, we did not analyze promoter-promoter chromatin contacts between duplicated genes, but focused instead on promoter-enhancer contacts. Indeed, we were mainly interested in understanding those features of the evolution of chromatin contacts that can be generalized outside of duplicated genes.

Finally, although we believe that many of the conclusions drawn from our analysis of duplicated genes hold true for the evolution of regulatory chromatin contacts in general, we emphasize that future evolutionary studies are needed to better investigate this important aspect of gene expression regulation. High-resolution chromatin contact datasets, comparable across species, are needed to establish a comprehensive and unbiased overview of the evolution of *cis*-regulatory landscapes.

## Materials and methods

### Genome annotation data

We downloaded genome annotation data for mouse (GRCm38.p6 genome assembly) from Ensembl v102 (Harrison et al. 2024) using BioMart (Smedley et al. 2009). The following fields were retrieved for each transcript: gene stable identifier, gene biotype, transcript stable identifier, transcript biotype, chromosome/scaffold name, gene start (bp), gene end (bp), strand, and transcription start site (TSS).

### PCHi-C data

In this study, we relied on the Promoter Capture Hi-C (PCHi-C) technique to define promoter-centered chromatin contacts. This technique is a derivative of Hi-C (Lieberman-Aiden et al. 2009) that specifically targets pre-defined gene promoters to enrich for putative regulatory chromatin contacts (Schoenfelder et al. 2015; Mifsud et al. 2015). The genome is divided into restriction fragments and interactions are scored between pairs of restriction fragments, including at least one targeted (baited) fragment. We use PCHi-C data that was processed by Laverré, Tannier, and Necsulea 2022, for mouse. These data were compiled from different sources and were analyzed with a homogeneous bioinformatic pipeline that uses HiCUP and ChICAGO (Wingett et al. 2015; Cairns et al. 2016). We provide the list of analyzed PCHi-C samples and their publication of origin in Table S1.

In all analyses, we used the coordinates of a set of mouse enhancers defined by the ENCODE consortium (Yue et al. 2014), which was also used by Laverré, Tannier, and Necsulea 2022. We downloaded the contacts between genes and this set of ENCODE-defined enhancers from the supplementary dataset 4 of Laverré, Tannier, and Necsulea 2022. We filtered these data to remove entries with missing information and retained only contacts occurring in *cis* (*i.e.* on the same chromosome). We removed contacts for which the genomic distance between gene TSS and enhancers was smaller than 25 kb or greater than 2 Mb.

### RNA-seq data processing

We aimed to analyze RNA-seq data and PCHi-C data for the same biological samples. We thus went back to the original publications describing PCHi-C data and retrieved SRA accession numbers for RNA-seq samples, when available. We were not able to retrieve RNA-seq data from two biological conditions, namely embryonic stem cells (ESC) and fetal liver cells (FLC) from Schoenfelder et al. 2015. All RNA-seq samples used here are publicly available and were downloaded from the SRA database (Kodama et al. 2012) using the prefetch and fastq-dump tools of the SRA Toolkit. Information on RNA-seq samples, including cell types, treatment / biological conditions and SRA accession numbers, is provided in Table S2.

Transcript expression levels were quantified with kallisto 0.51.1 (Bray et al. 2016). Transcript sequences (cDNA) were obtained from Ensembl (version 102), and the transcriptome was indexed with kallisto index. Transcript abundances were then estimated using kallisto quant, in unstranded mode and with default parameters. Among the resulting outputs, TPM (transcripts *per* million) values were retained. We used the R package tximport (Soneson, Love, and Robinson 2015) to compute gene-level TPMs, by summing the TPMs of all transcripts associated with each gene. Subsequently, the mean TPM across biological replicates was computed for each gene and for each cell type or biological condition. TPM values are typically computed using the following formula:

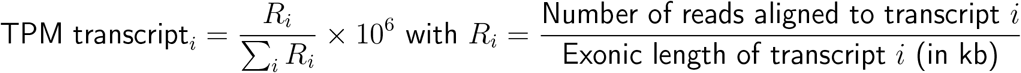

### Paralogous relationships

We downloaded data on paralogous gene pairs from Ensembl v102 using BioMart. The following fields were retrieved: stable identifiers for the two genes of each pair, last common ancestor, homology type, and the percent protein sequence identity for the two genes, computed using each gene in turn as the reference. We removed gene pairs annotated as “gene split” as they do not represent true paralogs.

As expected, some genes have numerous paralogs because they originate from gene families that have undergone multiple duplication events. To avoid giving too much weight to these families, we retained only one paralogous relationship *per* gene. Specifically, we kept the most recent duplication event for each gene, if and only if this event was not followed by subsequent duplications for one of the two gene copies in the same species. To do this, we downloaded the species tree and the gene family trees from Ensembl v102. Gene family trees were available in the MySQL database of Ensembl-Compara in a tabular format, where each entry corresponded to a branch in the tree. In this table, trees were classified as “tree“, “supertree” (which link different gene trees or subfamilies together when gene families are too large) and clusterset (which corresponds to a multiple analysis of trees that were not reconciliated). We discarded the “supertree” and the “clusterset” trees. We further retained only trees that were generated from protein sequences. We selected for further analysis only those trees that contained paralogous genes from our focal species. We wrote an R script to convert each tabular tree representation into a Newick format, for each gene family. Using functions from the ape package in R, we identified the most recent common ancestor (MRCA) for each pair of paralogs present in the gene trees. Finally, we retained paralogous pairs for which the clade descending from their MRCA did not contain additional paralogs from the same species. This corresponds to the filter named “terminal duplication nodes” in Fig. 1A.

We performed several filters on the PR dataset. First, we retained PRs for which both members were protein-coding. Second, we discarded PRs where one or both genes were assigned to unplaced contigs or to the mitochondrium. We also discarded PRs where the coordinates of the two genes were overlapping (as this suggests annotation errors rather than genuine duplication events). Third, we discarded PRs where there was a one-to-many exon difference, to remove potential retrotransposition duplication events. Fourth, as mentioned above, we retained PRs annotated as terminal duplication nodes. Fifth, we retained PRs where each gene was targeted by a single PCHi-C bait, and where the baits were different for the two genes. Moreover, we retained only those PRs for which contacts with ENCODE-defined enhancers were present.

We retrieved from Ensembl information on the phylogenetic tree node (last common ancestor) where the duplication event has likely occurred. We retrieved median divergence times from TimeTree (Kumar et al. 2022). We did not find divergence time information for the Murinae subfamily (this divergence time is represented as “NA” in Figure 1).

### Simulated datasets

We generated two types of control datasets for our analyses. The first, referred to as the “shuffled dataset”, was generated by shuffling the true paralogous gene pairs, creating new “paralogous” relationships within the same set of 2,840 genes. The second, referred to as the “sampled dataset”, mimics the characteristics of the true paralogous gene set, specifically with respect to genomic localization and expression levels. To construct this dataset, we first divided paralogous gene pairs into bins. Each bin was characterized by an expression level class and by a genomic localization class. Expression classes were created by dividing gene pairs into 20 equal-size classes, based on the average expression levels of the two genes in pair. The genomic localization class was defined as follows:

- for genes localized in *trans*, we defined the class as the pair of chromosomes on which the two genes were situated.
- for genes localized in *cis*, we defined the class as the combination of the chromosome on which the two genes were situated, and the distance between the two genes’ TSS. The distance was discretized by constructing 250 kb - wide distance bins (*i.e.* 0 - 250 kb, 250 - 500 kb *etc*).

We then constructed random gene pairs, drawn from the set of all protein-coding genes excluding our PR dataset, and we assigned them to the same expression-localization classes. Finally, we randomly drew sets of 1,420 protein-coding gene pairs that matched the distribution of the true PR dataset with respect to the expression-localization classes.

For each dataset type, we generated 1,000 replicates by varying the seed for random number generation from 1 to 1,000. For visualization purposes, one dataset out of the 1,000 replicates was selected for each control type, identifiable by the seed used to generate it. This selection was performed by computing the density curve of the median value of the analyzed metric across all sampled datasets and choosing the dataset whose median corresponded to the highest point of the density curve. The random seeds used to generate these typical datasets are provided in the figure legend.

### Gene Ontology enrichment

To test whether biological functions were different between young duplicates and old duplicates, we performed a Gene Ontology enrichment analysis using the GOrilla webserver (Eden et al. 2009). To do this, we first ordered the genes involved in the 1,420 PRs by their estimated duplication age. To identify functions enriched among younger duplicates, we ordered genes in ascending duplication age order. Conversely, to identify functions enriched among older duplicates, we ordered genes by descending duplication age, with older genes at the top of the list. The results of the enrichment analysis are provided in Table S3 and Table S4.

### Evaluating gene expression level predictions for paralogous genes

We used a simulation approach to test whether gene expression levels predicted with kallisto were correct for paralogous genes. Specifically, we simulated transcript-level and gene-level RNA-seq coverage values using yasim version 3.2.1 (Su et al. 2024). Using these values as input, we simulated Illumina-like RNA-seq reads, using art_modern version 1.3.2 (Yu 2026). We performed this analysis in single-end and paired-end modes, with a 75bp read length. The simulations were performed for all mouse genes and transcripts, starting from the Ensembl 102 annotation used everywhere in this manuscript. The simulation and evaluation procedure is detailed below.

First, to generate gene-level RNA-seq read coverage values, we used the generate_gene_-depth function in yasim, with the Ensembl 102 annotations in GTF format as an input. We asked for an average gene depth (or read coverage) of 10. Second, to generate transcript-level RNA-seq read coverage values, we used the generate_isoform_depth function of yasim, with the gene-level read coverage values obtained previously and the same gene annotation GTF file. Third, we used transcript-level read coverage values to generate 75bp Illumina-like reads, in both single-end (SE75) and paired-end (PE75) modes (see below). To do this, we used art_modern (Yu 2026) with the --mode trans and --read_len 75 options. We provided as an input the fasta file containing transcript cDNA sequences downloaded from Ensembl 102, as well as the transcript-level read coverage values obtained above. For the PE75 run, we set the average fragment length at 200 (--pe_frag_dist_mean 200) with a standard deviation of 20 (--pe_frag_dist_std_dev 20). We also used the --builtin_qual_file GA2Recalibrated_75bp option, to generate data with sequencing errors that are as similar as possible to the ones of real data. Finally, to evaluate gene expression levels on the simulated data, we used kallisto version 0.51.1 in the exact same way as was done for real data.

To estimate the expected number of reads and the expected TPM values for the simulated RNA-seq data, we used the following formulas:

For SE75 mode and a transcript *t*:

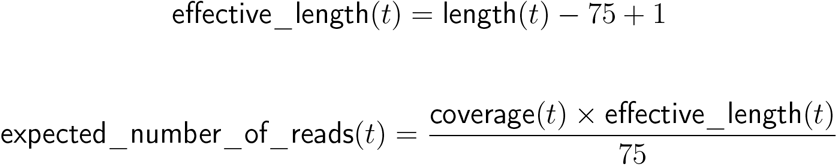

For PE75 mode, an average fragment length of 200 and a transcript *t*:

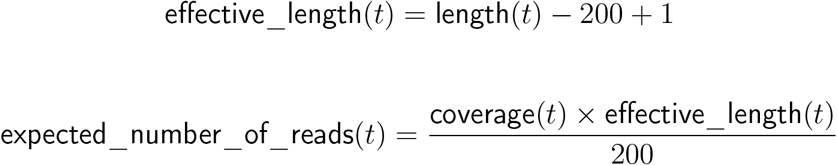

For both SE75 and PE75 modes, we then computed expected TPM values with the following formula:

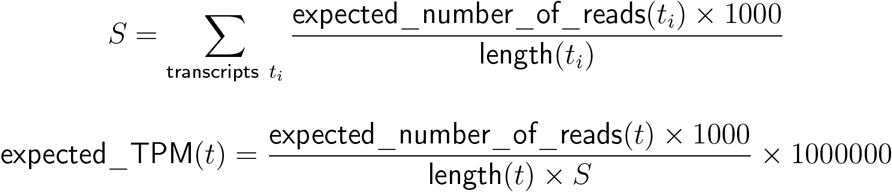

The TPM value of a gene were obtained by summing the TPM values of all its transcripts, for both expected and predicted values. Smooth scatterplots were used to represent the relationship between the expected and predicted gene TPM levels, for all 22,287 protein-coding genes and for the 2,840 paralogs. The same was done for the Δexpression values, computed on the expected and predicted gene expression levels, for the pairs of paralogs.

### Characterisation of the evolution of the expression of paralogous genes

We used two metrics to measure the expression divergence of paralogous gene pairs expression. The first metric, Δexpression, was calculated as the difference in overall expression levels (measured as the mean TPM across samples) between the two genes of each paralogous pair, normalized by the average expression level of the two genes:

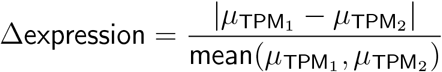

Δexpression values range from 0 to 2, where 0 indicates identical mean expression between the genes and 2 corresponds to one gene being expressed while the other is silent.

We then aimed to measure the difference between expression profiles across biological samples. To do this, we first calculated relative TPMs as described in Laverré, Tannier, and Necsulea 2022, to reduce the effects of scale differences between genes. Relative TPMs were computed as follows:

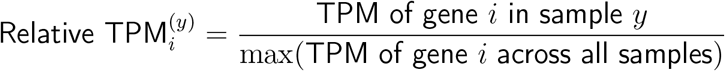

The second metric, referred to as the “expression profile divergence” was computed using the Manhattan distance between the relative TPM vectors of each paralog pair:

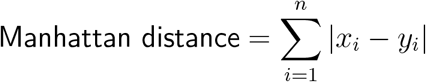

where *x* and *y* correspond to the relative TPM values of a paralogous gene pair across samples, and *i* indexes the samples. High Manhattan distance values indicate important differences in the expression profiles of the two paralogs. Conversely, low Manhattan distance values indicate that the two expression profiles are similar.

### Characterisation of the evolution of promoter-enhancer contacts

In this study, we defined the regulatory complexity of a gene as the number of unique enhancers in contact with this gene in the PCHi-C data. For each gene pair, we defined Δcomplexity as follows:

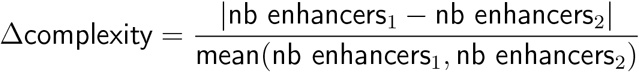

Thus, Δcomplexity measures the normalized difference in chromatin contact complexity between the two gene pairs.

We then aimed to assess the difference between chromatin contact profiles across biological samples. To do this, we first counted the number of enhancers contacted by each gene in each sample, for all genes that were targeted in the PCHi-C data (not only those involved in paralogous relationships). This value was then divided by the total number of gene–enhancer contacts in each sample, across all genes, and multiplied by one million. We thus obtained a value that we name EPM for “enhancers *per* million”, which is analogous to TPM:

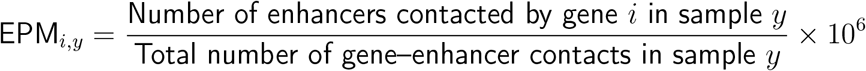

We then computed relative EPMs across samples, as follows:

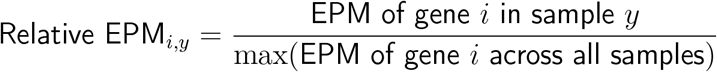

The divergence between chromatin contact profiles of paralog pairs was then measured using the Manhattan distance:

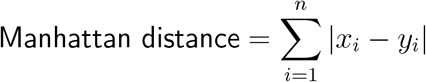

where *x* and *y* correspond to the relative EPM values of a paralogous gene pair across samples, and *i* indexes the samples.

### Coding sequence evolution

To evaluate coding sequence evolution, we computed the ratio of non-synonymous substitutions *per* non-synonymous site (dN) and the ratio of synonymous substitutions *per* synonymous site (dS). To do this, we extracted the longest transcript for each gene and we aligned the corresponding coding sequences for each pair of paralogous genes using PRANK (Löytynoja 2014) in codon mode. We then computed dN and dS values for the gene pair with the codeml program within PAML version 4.9j (Yang 1997), setting the cleandata parameter at 0 (meaning that sites with ambiguity data are kept) and the model parameter at 0 (meaning that all branches are assumed to have the same dN/dS ratio). We used default values for all other parameters.

### Promoter sequence evolution

To define gene promoters, we first retrieved APPRIS annotation scores (Rodriguez et al. 2022) for each protein-coding transcript, using the Ensembl BioMart interface (Harrison et al. 2024; Smedley et al. 2009). These scores provide a means to select functionally important isoforms. We ranked the transcripts in decreasing order of predicted functional importance, from principal1 to principal5 and from alternative1 to alternative5. For each gene, we selected the transcript with the best APPRIS score, which we considered to be the “major isoform”. We then defined the promoter for the major isoform as the 1 kb region situated directly upstream of the transcription start site. We thus considered a single promoter region for each gene.

To evaluate promoter sequence evolution, we extracted DNA sequences containing the entire gene regions and a 10 kb flanking region on each side, for each paralogous gene pair. We aligned these pairs of genomic sequences using Progressive Cactus version 2.9.1 (Armstrong et al. 2020). We transformed these alignments from HAL to MAF format using the cactus-hal2maf tool provided with Progressive Cactus. We then queried these alignments with the mafsInRegion tool provided by the UCSC GenomeBrowser, to retrieve alignments corresponding to each gene’s promoter region. We then evaluated the percentage of aligned sites and the percentage of identical sites with respect to the promoter size, for each aligned promoter. We obtained two values, one using as a reference the promoter of the first gene in the pair, and one using as a reference the promoter of the second gene in the pair. We used the average of the two values in our analyses. We note that this approach does not require that the two promoters be homologous. For each gene promoter, we analyze the sequence that it aligns with, in the neighbourhood of the paralogous gene, but we do not require that this sequence correspond to the promoter of the paralogous gene.

As we found that promoter sequences are not well aligned between old paralogous genes, we defined an additional measure of promoter sequence similarity based on the presence of shared transcription factor binding sites. Specifically, we used the FIMO tool from the MEME suite (Bailey et al. 2009), version 5.5.7, to find transcription factor binding motifs in promoter sequences, defined as above. We used the JASPAR 2026 database (non-redundant vertebrate motifs) as a source for transcription factor binding motifs (Ovek et al. 2026). We ran FIMO with a *p*-value threshold of 10*^−^*^4^, setting the “maximum stored scores” parameter at 50 *×* 10^6^ to avoid changes in stringency at runtime. For each pair of promoter sequences (from a pair of paralogous genes or from a random gene pair), we computed a Jaccard index of the sets of transcription factors that had at least one FIMO-identified motif occurrence in each promoter.

### Common contacted enhancers

Having observed that *cis*-localized duplicated genes have higher levels of chromatin contact similarity than *trans*-localized duplicated genes, we wanted to verify if part of this observation can be explained by enhancer sharing. Indeed, paralogs lying on the same chromosome could contact the same enhancers. To see how often this is the case and whether this is more than expected by chance, we quantified the proportion of shared enhancers between two *cis*-localized using the Jaccard index. Specifically, for *cis*-localized gene pairs found within a maximum distance of 4 Mb (computed between the gene TSS), we defined the following sets:

- the set of enhancers that are contacted by at least one gene and that can be contacted by both genes. This corresponds to the union of the sets of enhancers that are contacted by at least one of the two genes, and which are found at a distance between 25 kb and 2 Mb with respect to the TSS of both two genes. This corresponds to a modified union of the sets of enhancers contacted by the two genes.
- the set of enhancers that are contacted by both genes. This corresponds to the intersection of the sets of enhancers contacted by the two genes.

Here, promoter-enhancer contacts are considered across all PCHi-C samples combined. We defined a Jaccard index as the ratio between the number of enhancers in the second set and the number of enhancers in the first set. This process was applied to the PRs and to the 1000 “sampled” control sets which have the same proportions of *cis*-localized pairs as the PRs.

### Ohnolog dataset

We downloaded the annotation of paralogous genes originating from the 2 whole-genome duplications in the ancestor of all vertebrates (2R-WGD) known as *ohnologs* from the OHNOLOGS.V2 database (Singh and Isambert 2020). We selected the strictest available annotation, defined by the following criteria: q-score (outgroup) < 0.001 and q-score (self) < 0.001. The q-score is a database-defined metric ranging from 0 to 1 that estimates the probability that an *ohnolog* pair is identified by chance.

These annotations were then intersected with our paralog dataset (1,420 pairs), and only pairs annotated as both *trans* and *very old* paralogous pairs were retained as being an ohnolog pair. This filtering was applied because we considered ohnologs located in *cis*, as well as those corresponding to duplication events occurring after vertebrate speciation according to Ensembl, to be the result of technical annotation errors.

### Statistics and graphical representations

All statistical analyses and graphical representations were done in R-4.5.3 (R Core Team 2024). We used non-parametric statistical tests to perform comparisons. We tested the normality of the expression profile divergence and the chromatin contact profile divergence using the Shapiro-Wilks test in R. We used linear regression models to analyze relationships among these variables and additional factors (including expression levels and genomic localization).

We used smooth scatter plots to represent relationships between two continuous variables, using the smoothScatter function from the graphics package in R. This function uses a 2D kernel density estimate to produce a color density representation of a scatter plot. We used functions in the vioplot (Adler et al. 2025) package in R to display violin plots. These plots combine boxplots and density plots to represent the distribution of a continuous variable.

## Data availability

All processed data generated in this study are available online in a Zenodo repository (https://doi.org/10.5281/zenodo.21392106). All scripts used in this analysis are available on GitLab https://gitlab.in2p3.fr/victor.lefebvre/duplicon.

## Acknowledgements

We would like to thank Marie Sémon for her helpful suggestions on the manuscript. This work was performed using the computing facilities of the CC LBBE/PRABI. We are also grateful to the genotoul bioinformatics platform Toulouse Occitanie for providing computing and storage resources. V. Lefebvre was supported by a PhD fellowship from the French Ministry of Higher Education and Research.

