## SupplementaryFiguresAndTables for "Evolution of regulatory chromatin contacts: insights from duplicated genes"

#### Supplementary Material

### Supplementary Figures

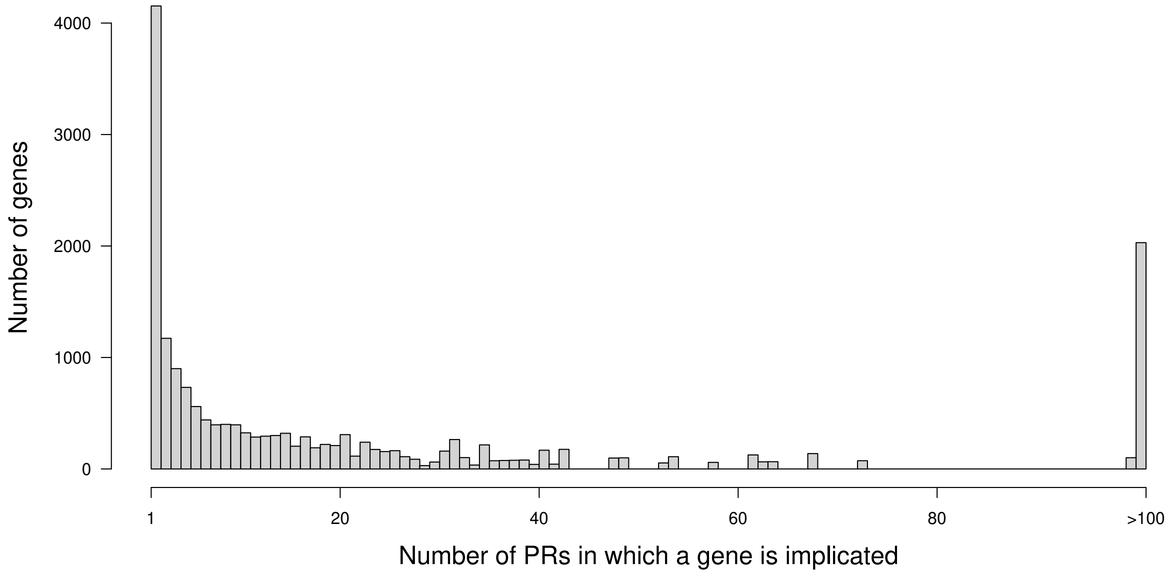


**Figure S1:** Histogram showing the distribution of the number of paralogous relationships (PRs) *per* gene. This analysis was performed on the entire set of PRs involving protein-coding genes, from the Ensembl Compara database, for the mouse genome.

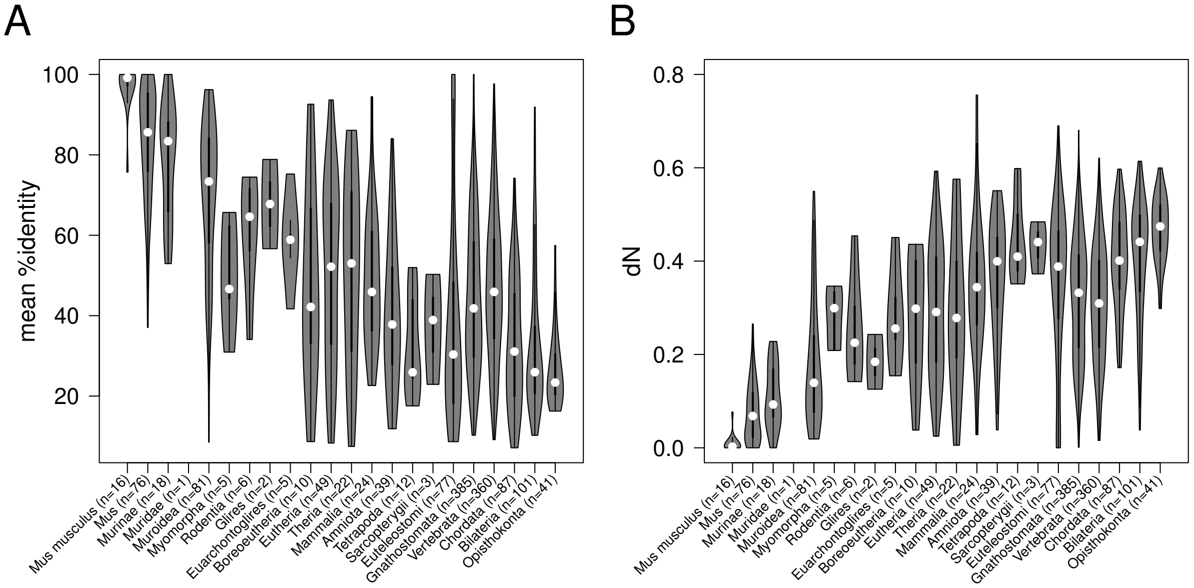


**Figure S2:** Relationship between protein sequence divergence and duplication age. **A.** Violin plots representing the distribution of the percentage protein sequence identity for paralogous gene pairs, as a function of duplication age. **B.** Violin plots representing the distribution of the rate of non-synonymous substitutions (dN) for paralogous gene pairs, as a function of duplication age.

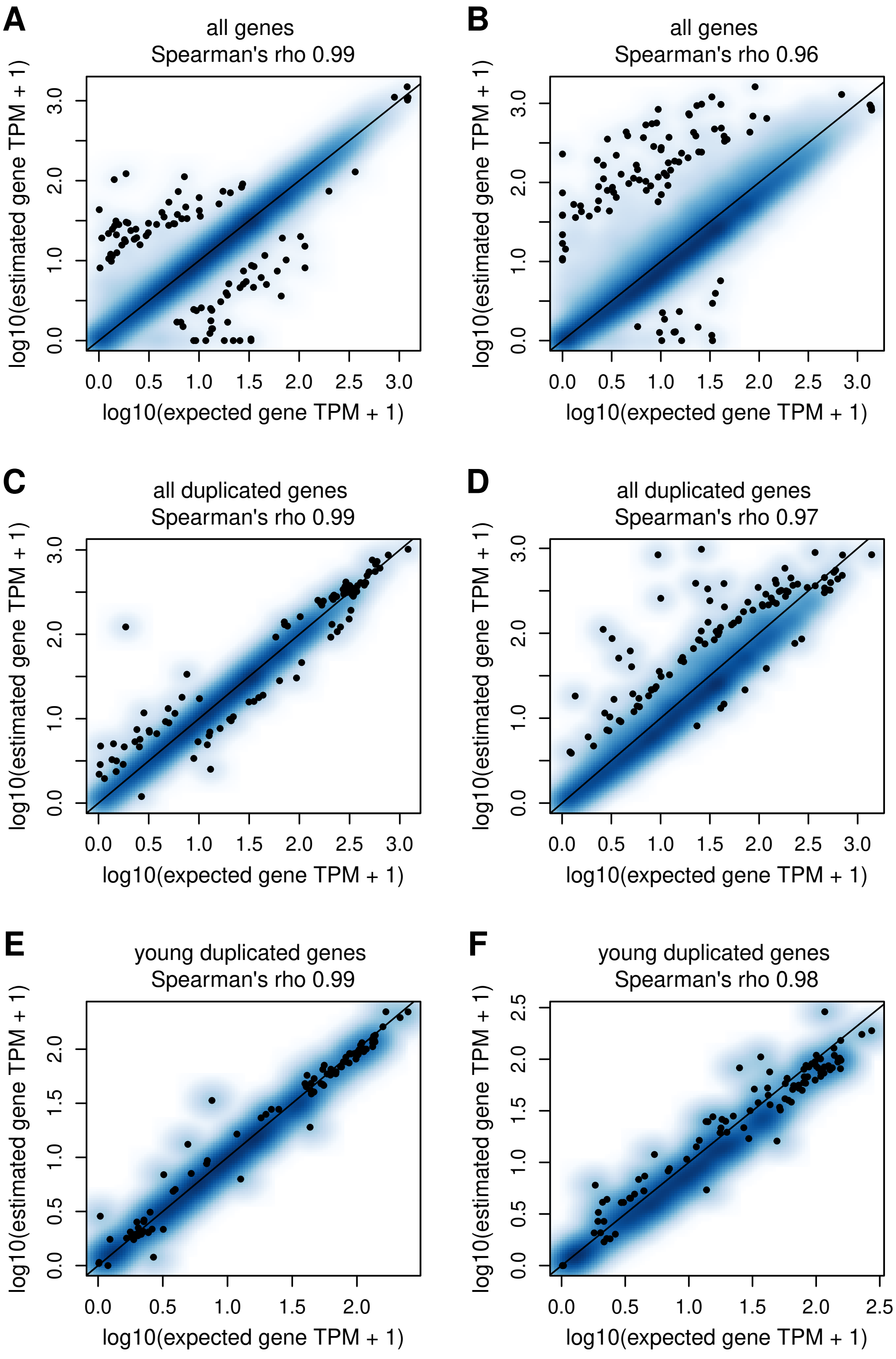


**Figure S3:** `Kallisto` predicts well gene expression levels for paralogous genes. **A**. Smooth scatterplot representing the relationship between the expected gene expression level and the `Kallisto`-predicted gene expression levels (log-transformed TPM), for all protein-coding genes, in SE75 read simulation mode. **B.** Same as **A**, in PE75 read simulation mode. **C.** Same as **A**, for the 2,840 paralogous genes included in our study, in SE75 mode. **D.** Same as **C**, in PE75 read simulation mode. **E.** Same as **A**, for the 410 paralogous genes belonging to "young" PRs. **F.** Same as **E**, in PE75 read simulation mode.

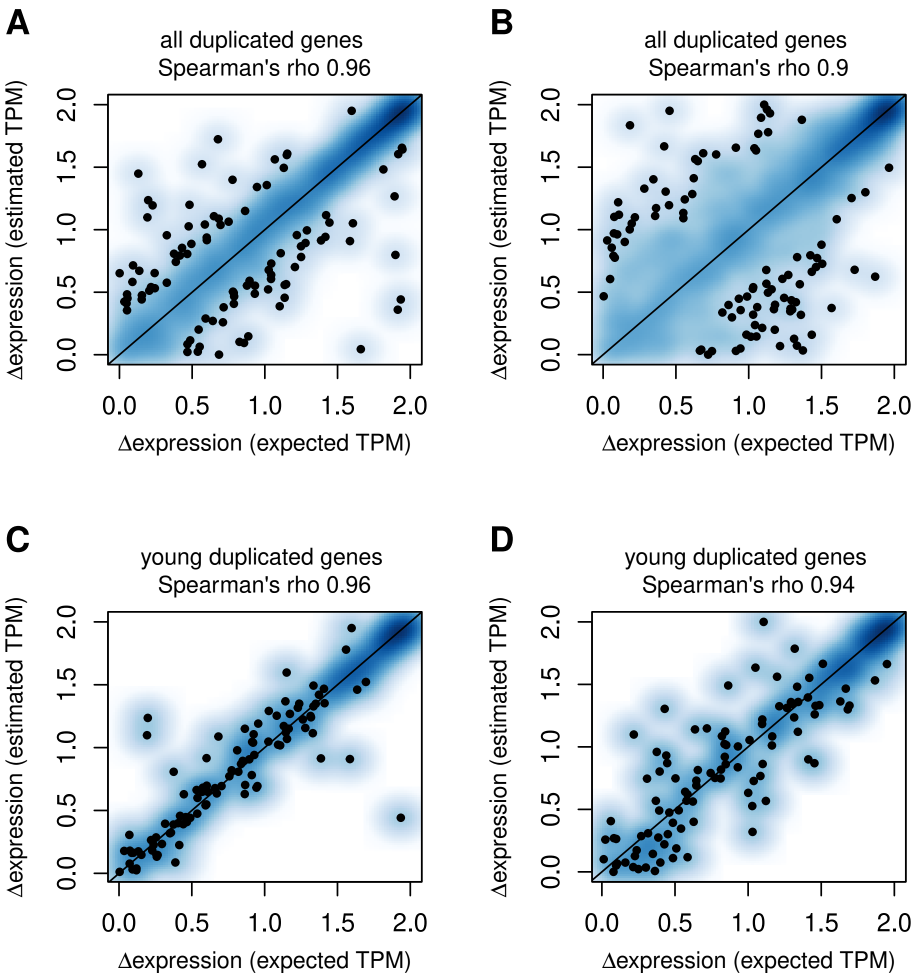


**Figure S4:** `Kallisto` predicts well gene expression levels for paralogous genes. **A.** Smooth scatterplot representing the relationship between the expected and estimated Δexpression values for the 1,420 PRs, in SE75 simulation mode. **B.** Same as **A**, in PE75 simulation mode. **C.** Same as **A**, for the 205 PRs belonging to the "young" age class. **D.** Same as **C**, in PE75 simulation mode.

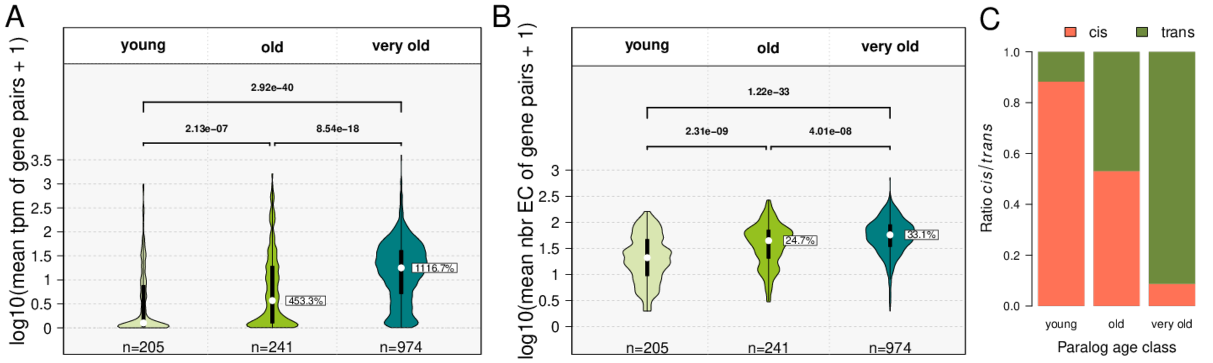


**Figure S5:** **A.** Violin plots showing the distribution of the average expression levels of duplicated genes, according to the approximate age of their duplication event: *young* corresponds to duplications that occurred between *Mus musculus* and Glires, *old* to duplications from *Euarchontoglires* to *Teleostomi*, and *very old* to duplications older than *Gnathostomata*. **B.** Same as **A**, but for the number of contacted enhancers. **C.** Barplot showing the ratio of *cis* and *trans* paralogous gene pairs according to their age classes. For panels **A**–**B**, *p*-values correspond to Wilcoxon tests; bold *p*-values indicate statistical significance. White rectangles indicate the relative difference between the medians (white dots) of the leftmost violin and the labeled violin.

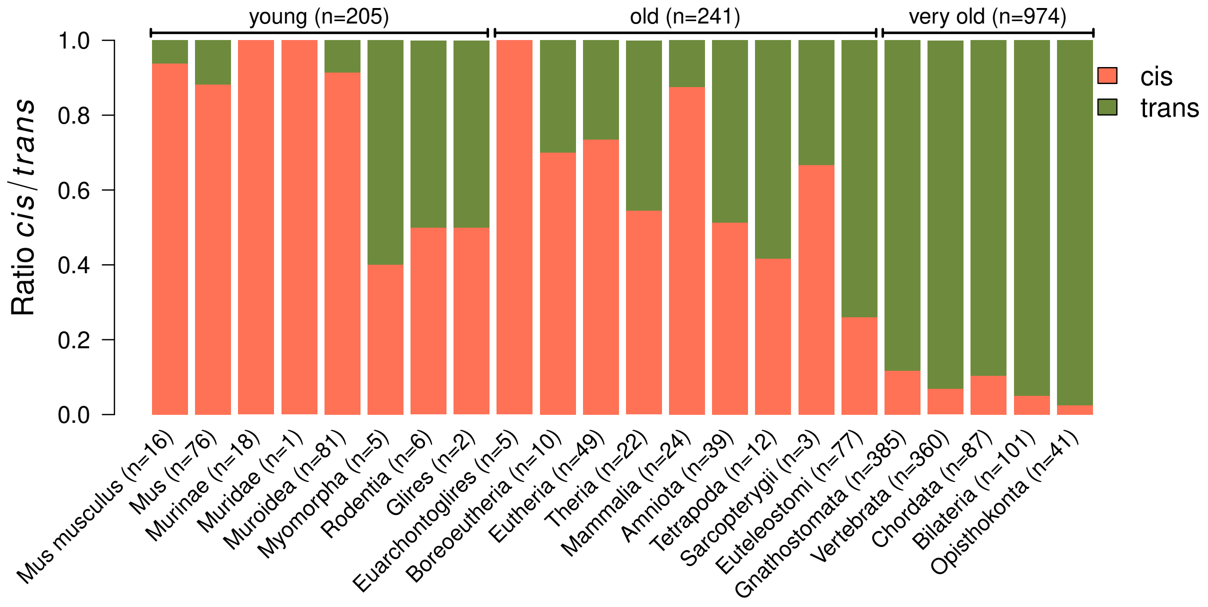


**Figure S6:** Proportion of paralogous pairs located in *trans* (green) and in *cis* (orange), according to the age of their duplication event.

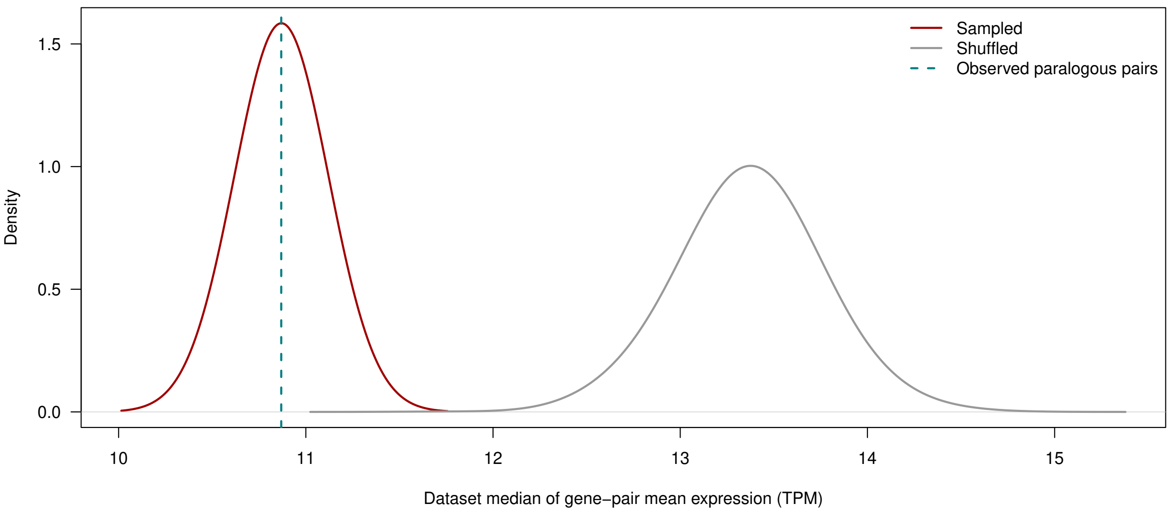


**Figure S7:** For each of the 1,000 sampled control datasets (red) and 1,000 shuffled control datasets (grey), the mean expression level was first calculated for each gene pair, and the median of the median of these pairwise means values was then calculated across the dataset. Density curves represent the distributions of the resulting 1,000 medians. The dashed line indicates the corresponding median calculated for the observed paralogous pairs

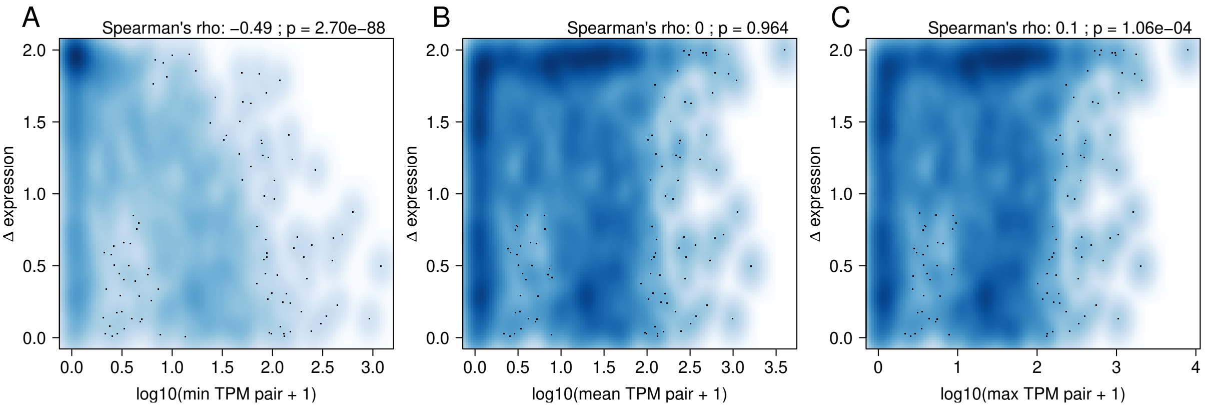


**Figure S8:** Correlation between Δexpression and expression levels of paralogous genes. **A.** Smooth scatter plot representing the relationship between Δexpression and the minimum expression level (log-transformed) of the two genes in the paralog pair, across the 1,420 paralogous gene pairs. **B.** Same as **A**, for the relationship between Δexpression and the mean expression level of the two genes in the paralogous pair. **C.** Same as **B**, for the relationship between Δexpression and the maximum expression level of the two genes in the paralogous pair.

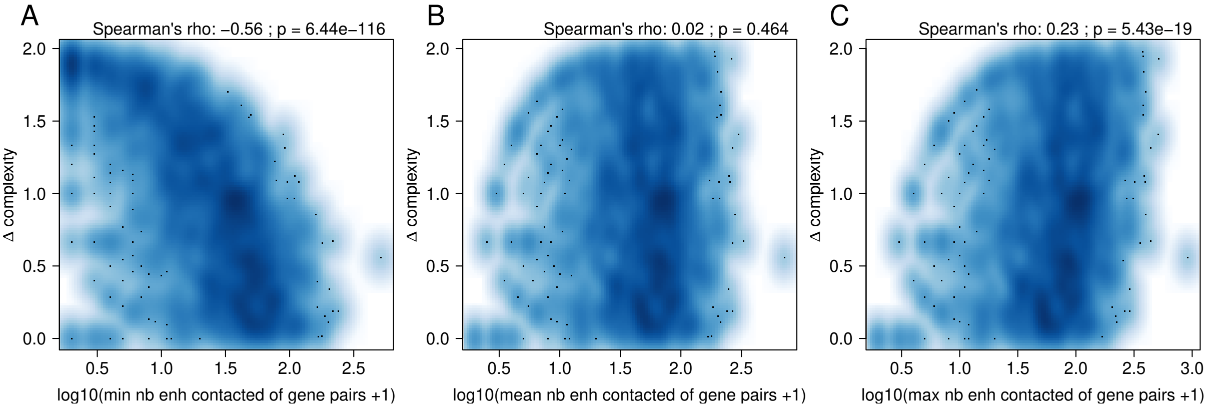


**Figure S9:** Correlation between $\Delta$complexity and the number of enhancers contacted by paralogous genes. **A.** Smooth scatter plot representing the relationship between $\Delta$complexity and the minimum number of contacted enhancers by the two genes in the paralogous pair (log-transformed), across the 1,420 paralogous gene pairs. **B.** Same as **A**, for the relationship between $\Delta$complexity and the mean number of contacted enhancers by the two genes in the paralogous pair. **C.** Same as **B**, for the relationship between $\Delta$complexity and the maximum number of contacted enhancers by the two genes in the paralogous pair.

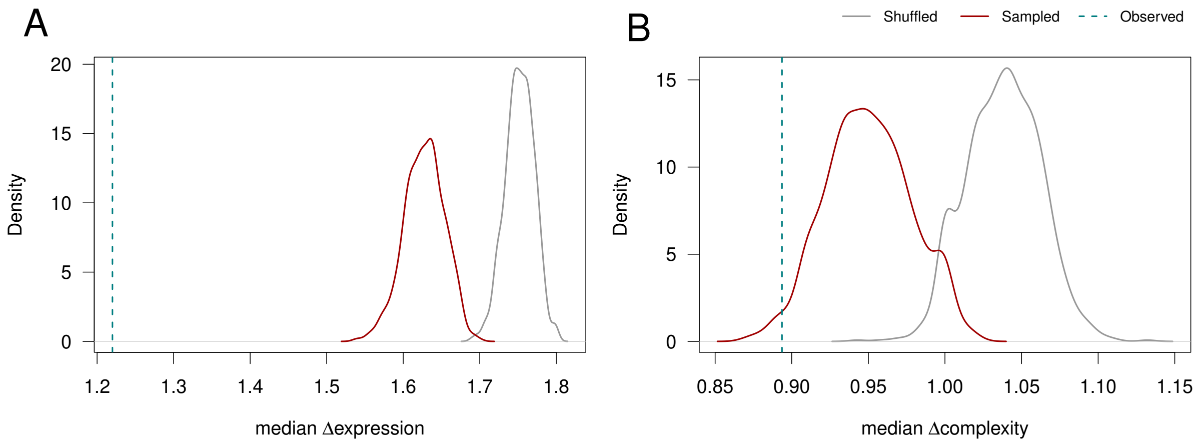


**Figure S10:** Distribution of the Δcomplexity and of the Δexpression across simulated datasets. **A**. Density plot representing the distribution of the median Δexpression across 1000 sampled datasets (red) and 1000 shuffled datasets (gray). The dotted vertical line represents the observed median value for PRs. **B.** Same as **A**, for the median Δcomplexity.

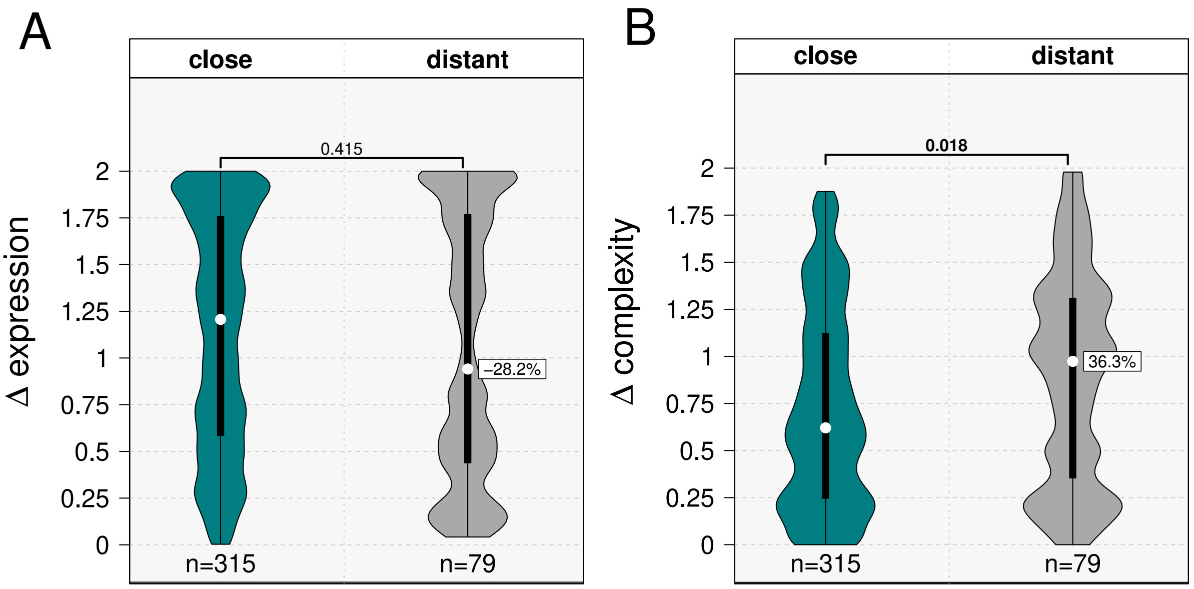


**Figure S11: A.** Violin plots showing the distribution of Δexpression values for *cis*-localized PRs, divided into two classes depending on the genomic distance between the two gene transcription start sites (TSS): the "close" class regroups genes with a between-TSS genomic distance below 1 Mb, the "distal" class regroups genes with a between-TSS genomic distance above 1 Mb. **B.** Same as **A**, for Δcomplexity. For panels **A**–**B**, *p*-values correspond to Wilcoxon tests; bold *p*-values indicate statistical significance. White rectangles indicate the relative difference between the medians (white dots) of the leftmost violin and the labeled violin.

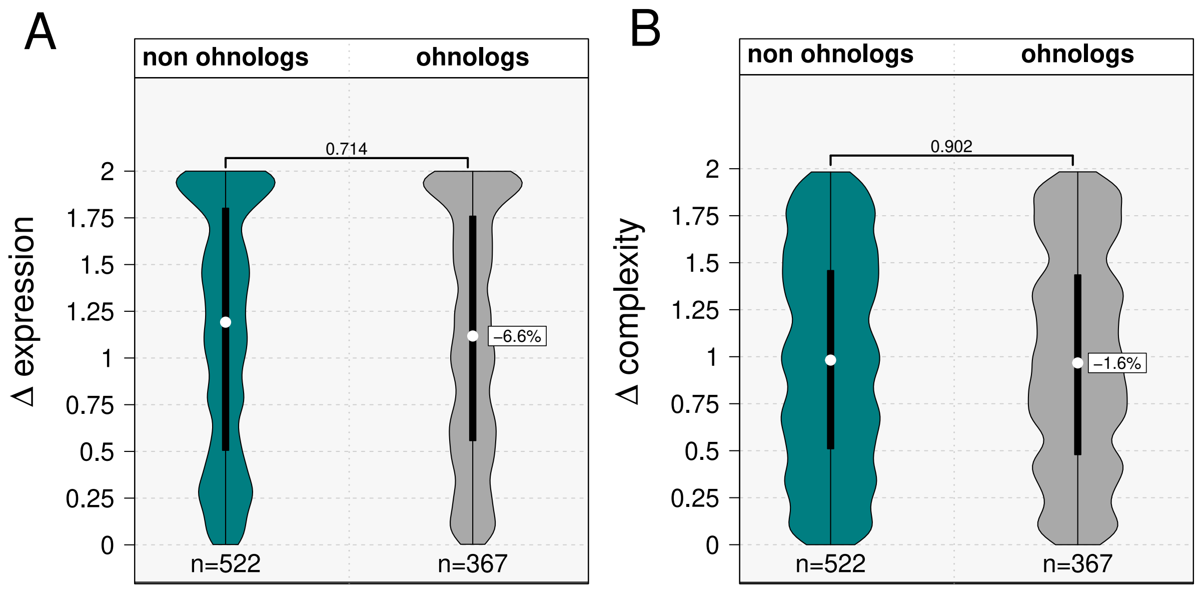


**Figure S12:** **A.** Violin plots showing Δexpression for *trans* paralog pairs separated in *ohnologs* and *non ohnologs* pairs. **B.** Same as **A** but for Δ complexity. For panels **A**–**B**, *p*-values correspond to Wilcoxon tests; bold *p*-values indicate statistical significance. White rectangles indicate the relative difference between the medians (white dots) of the leftmost violin and the labeled violin.

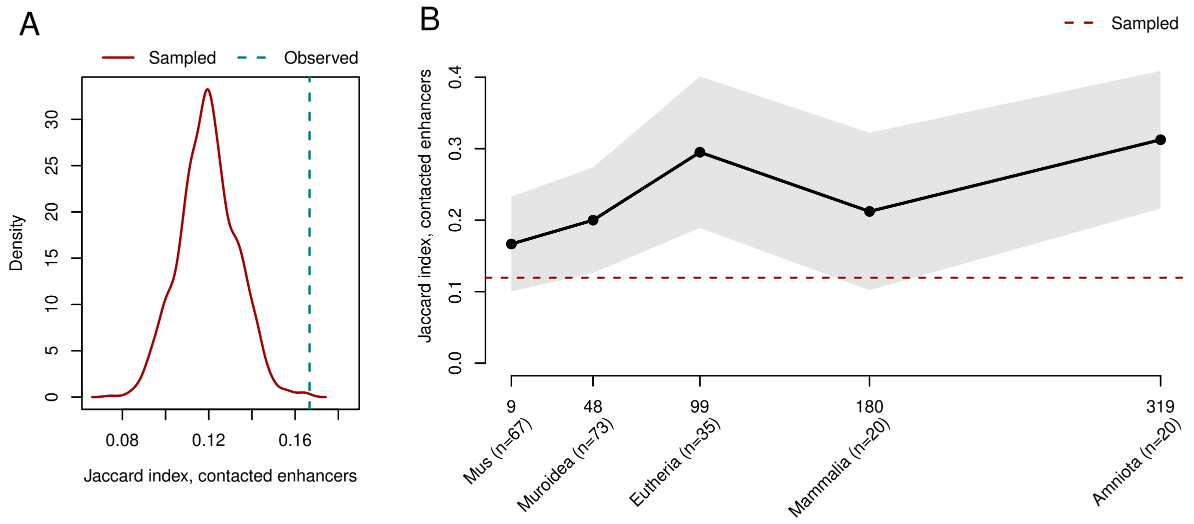


**Figure S13:** **A.** Density plot representing the distribution of the median proportion of common contacted enhancers (computed as the Jaccard index of the sets of enhancers contacted by the two genes, Materials and methods) for *cis*-localized gene pairs, for 1000 sampled datasets. The vertical bar represents the median proportion of common enhancers for *cis*-localized paralogous gene pairs. **B.** Distribution of the median proportion of common contacted enhancers, for *cis*-localized paralogous gene pairs, as a function of the duplication age.

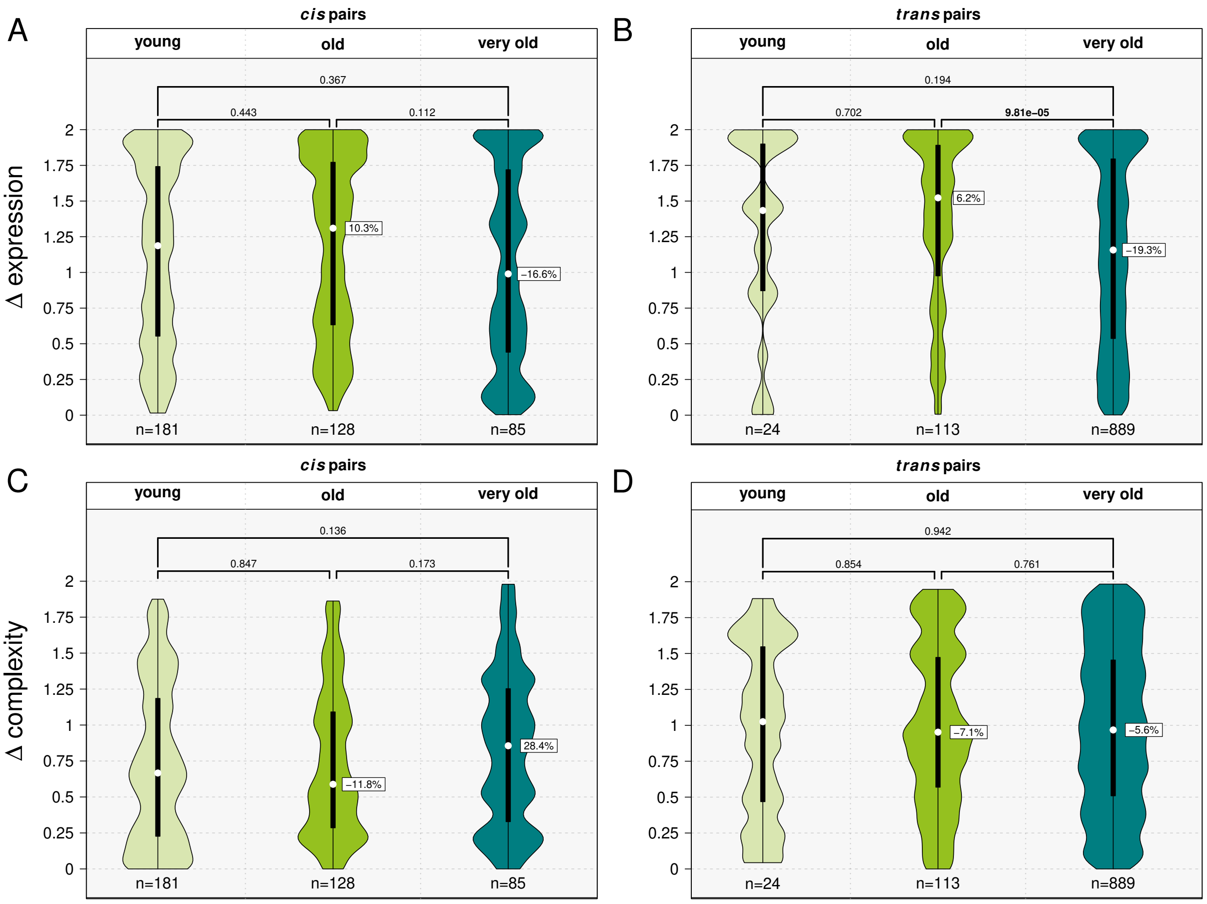


**Figure S14: A.** Violin plots showing the distribution of Δexpression values for *cis* paralog pairs, grouped according to the approximate age of their duplication event: *young* corresponds to duplications that occurred from *Mus musculus* to the Glires, *old* from *Euarchontoglires* to *Teleostomi*, and *very old* to duplications older than *Gnathostomata*. **B.** Same as **A** for *trans* paralog pairs. **C–D.** Same as **A–B**, but for Δ complexity. For panels **A–D**, *p*-values correspond to Wilcoxon test; bold *p*-values indicate statistical significance. The numbers indicated in white rectangles indicate the relative difference between the medians (white dots) of the leftmost violin and the labeled violin.

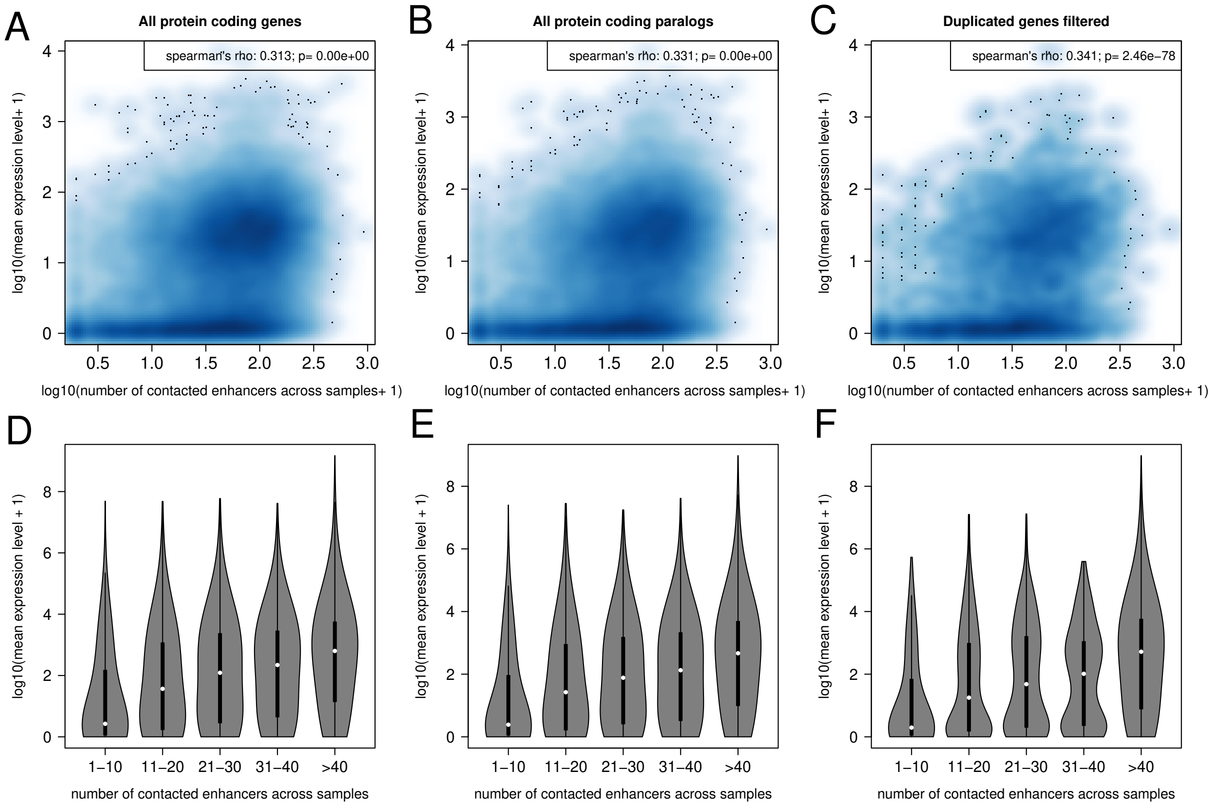


**Figure S15:** Correlation between expression levels and regulatory complexity. **A.** Smooth scatter plot representing the relationship between gene expression levels (averaged across all analysed RNA-seq samples) and regulatory complexity (total number of contacted enhancers) for all protein-coding genes. **B.** Same as **A**, for all protein-coding genes that are part of paralogous relationships. **C.** Same as **A**, for the 2,840 genes included in the final set of paralogous relationships analysed in this study. **D.** Violin plots representing the distribution of the average gene expression level. Genes were divided into 5 groups depending on the total numbers of contacted enhancers. All protein-coding genes are included in the analysis. **E.** Same as **D**, for all protein-coding genes that are part of paralogous relationships. **C.** Same as **D**, for the 2,840 genes included in the final set of paralogous relationships analysed in this study. For correlations with p-values reported as 0, values are below the numerical detection limit of R.

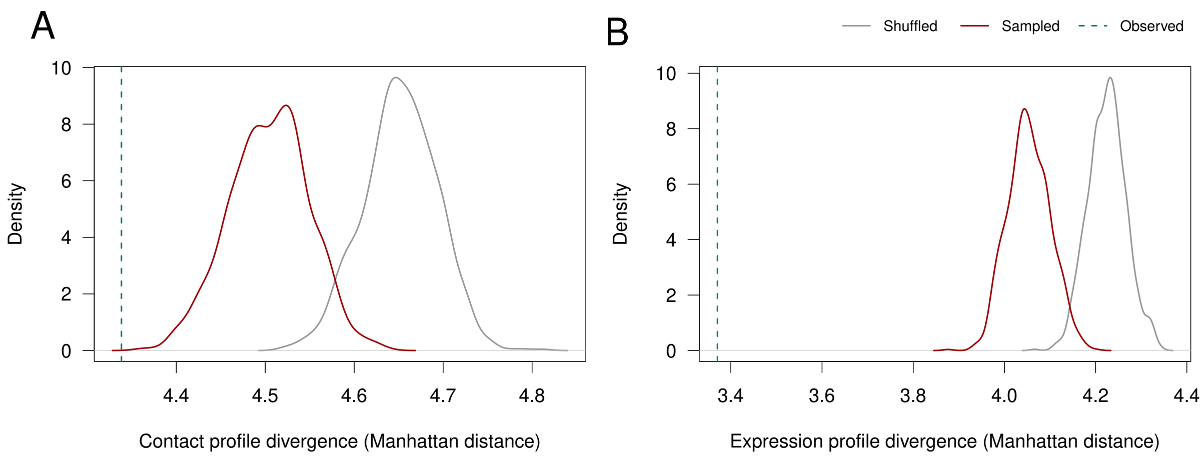


**Figure S16:** Distribution of the contact profile divergence and of the expresion profile divergence across simulated datasets. **A**. Density plot representing the distribution of the median contact profile divergence across 1000 sampled datasets (red) and 1000 shuffled datasets (gray). The dotted vertical line represents the observed median value for PRs. **B.** Same as **A**, for the median expression profile divergence.

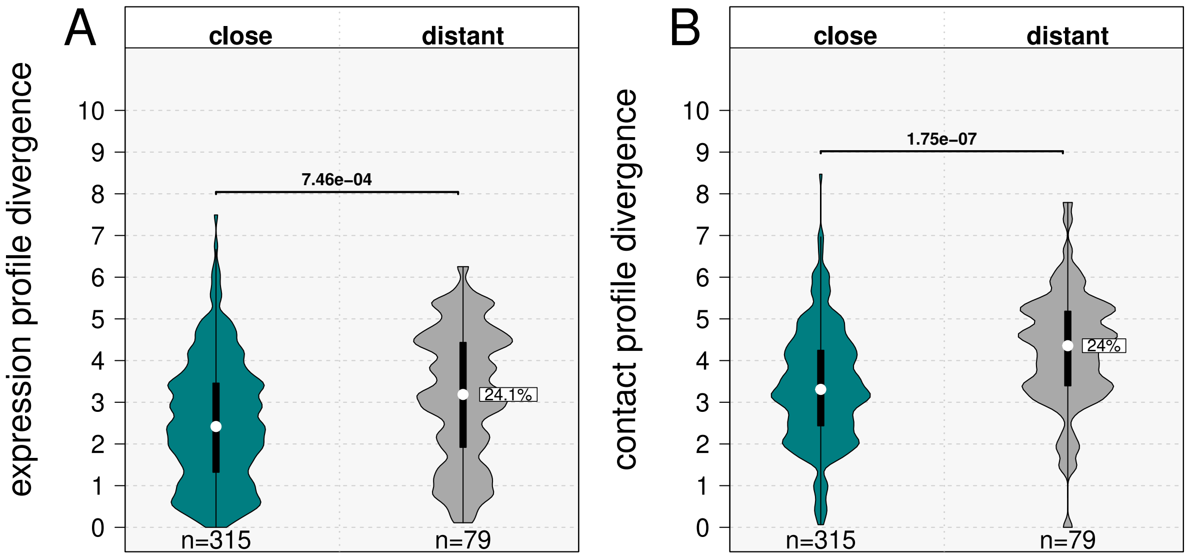


**Figure S17:** Relationship between expression profile divergence, contact profile divergence and genomic localization. **A.** Violin plot representing the distribution of the expression profile divergence for *cis*-localized genes, found in close distance to each other (maximum distance 1Mb, blue-green), or distant to each other (minimum distance 1Mb, gray). **B.** Same as **A.**, for the contact profile divergence.

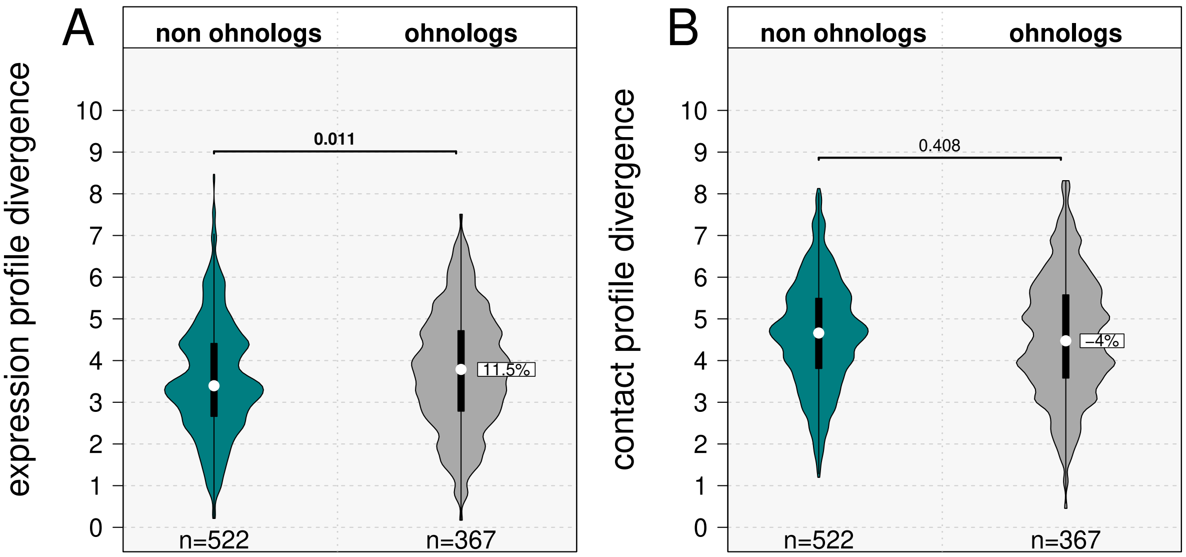


**Figure S18: A.** Violin plots showing the expression profile divergence for *trans* paralog pairs separated in *ohnologs* and *non ohnologs* pairs. **B.** Same as **A** but for the contact profile divergence. For panels **A**–**B**, *p*-values correspond to Wilcoxon tests; bold *p*-values indicate statistical significance. White rectangles indicate the relative difference between the medians (white dots) of the leftmost violin and the labeled violin.

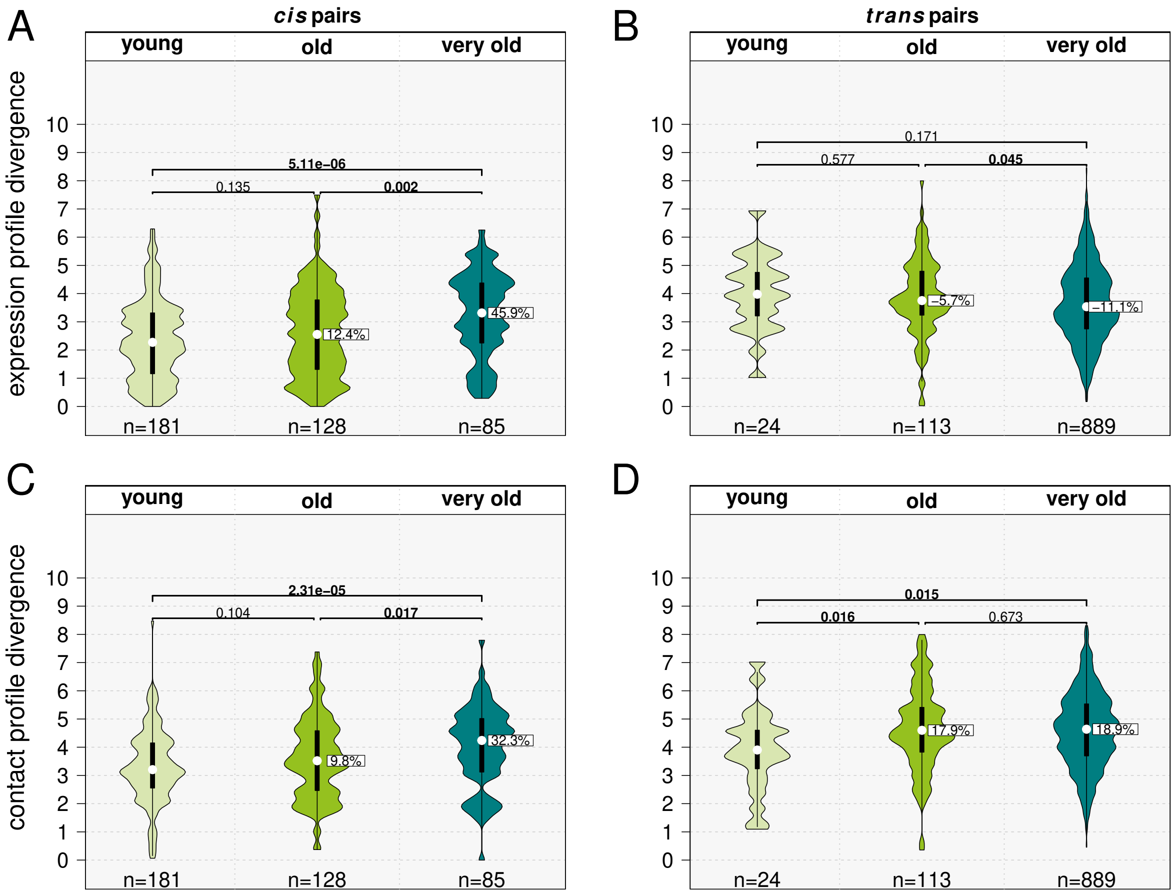


**Figure S19:** **A.** Violin plots showing expression profile divergence for *cis* paralog pairs grouped according to the approximate age of their duplication event: *young* corresponds to duplications that occurred from *Mus musculus* to the Glires, *old* from *Euarchontoglires* to *Teleostomi*, and *very old* to duplications older than *Gnathostomata*. **B.** Same as **A**, for *trans* paralog pairs. **C–D.** Same as **A–B**, but for contact profile divergence. For panels **A–D**, *p*-values correspond to Wilcoxon tests; bold *p*-values indicate statistical significance. White rectangles indicate the relative difference between the medians (white dots) of the leftmost violin and the labelled violin.

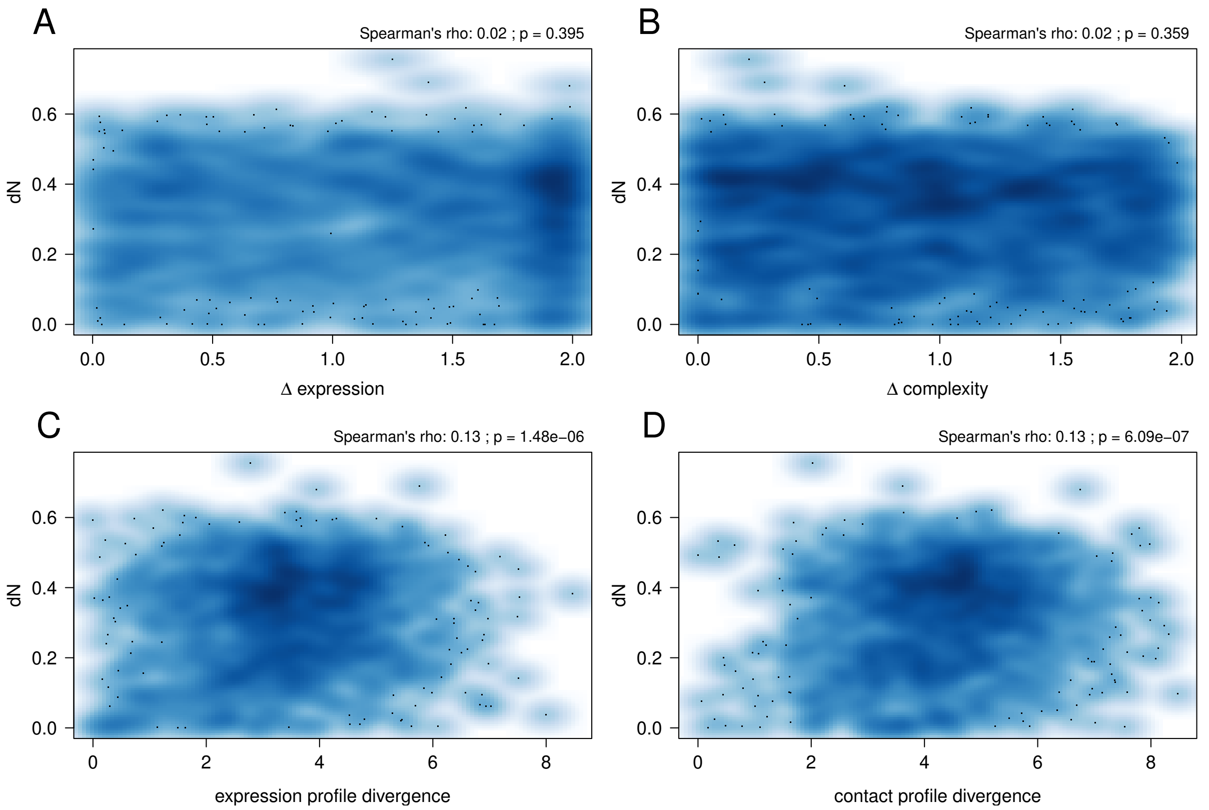


**Figure S20:** Correlation between expression and regulatory divergence metrics, on one hand, and protein sequence evolution, on the other hand. **A.** Scatter plot representing the relationship between Δexpression (on the X-axis) and the non-synonymous substitution rate (dN, on the Y-axis). **B.** Same as **A**, for Δcomplexity and dN. **C.** Same as **A**, for the expression profile divergence and dN. **D.** Same as **A**, for the contact profile divergence.

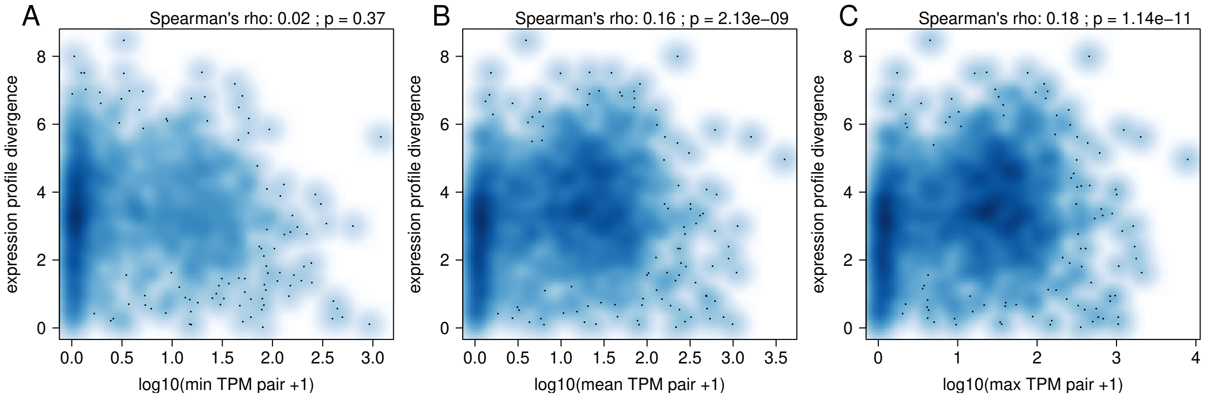


**Figure S21:** Correlation between the expression profile divergence and the expression levels of paralogous genes. **A.** Smooth scatter plot representing the relationship between the expression profile divergence and the minimum expression level (log-transformed) of the two genes in the paralog pair, across the 1,420 paralogous gene pairs. **B.** Same as **A**, for the relationship between the expression profile divergence and the mean expression level of the two genes in the paralogous pair. **C.** Same as **B**, for the relationship between the expression profile divergence and the maximum expression level of the two genes in the paralogous pair.

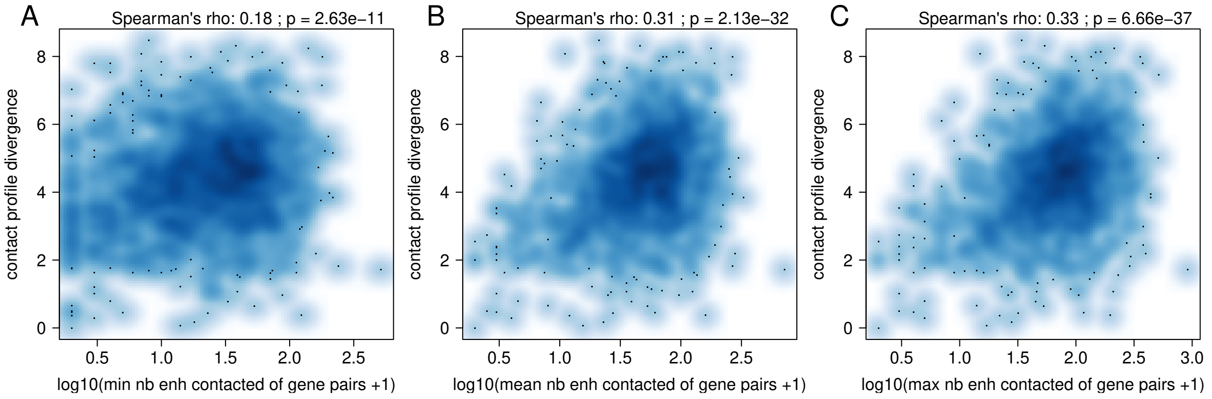


**Figure S22:** Correlation between the contact profile divergence and the number of enhancers contacted by paralogous genes. **A.** Smooth scatter plot representing the relationship between the contact profile divergence and the minimum number of contacted enhancers by the two genes in the paralogous pair (log-transformed), across the 1,420 paralogous gene pairs. **B.** Same as **A**, for the relationship between the contact profile divergence and the mean number of contacted enhancers by the two genes in the paralogous pair. **C.** Same as **B**, for the relationship between the contact profile divergence and the maximum number of contacted enhancers by the two genes in the paralogous pair.

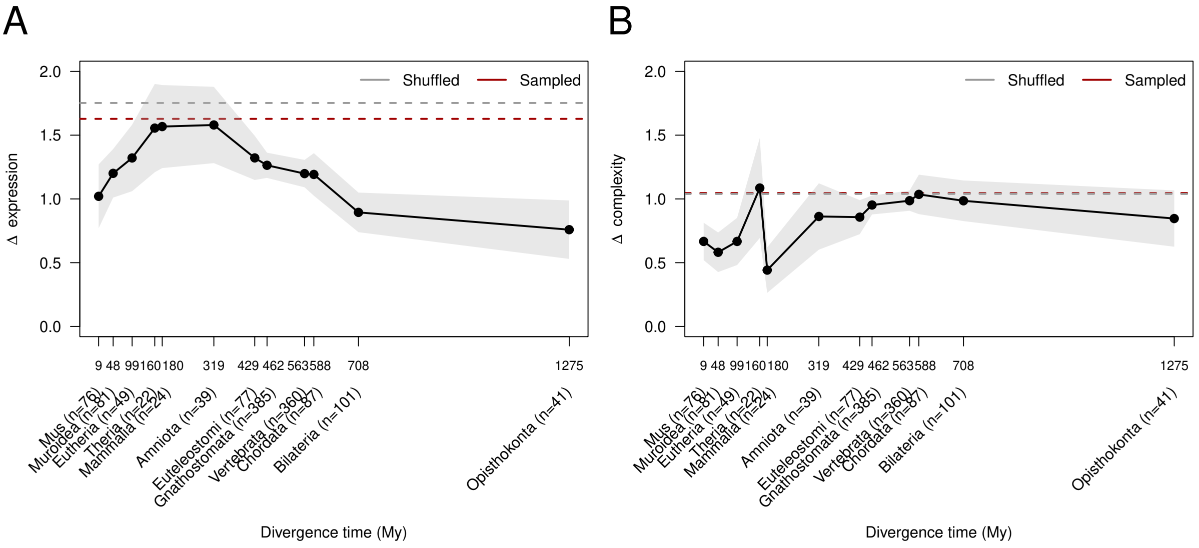


**Figure S23:** Dynamics of the expression level divergence and regulatory complexity divergence with time. **A.** Distribution of the median $\Delta$expression as a function of the estimated divergence time of paralogous gene pairs. Sampled median value: 1.63. Shuffled median value: 1.75 **B.** Distribution of the median $\Delta$complexity as a function of the estimated divergence time of paralogous gene pairs. Sampled median value: 1.046. Shuffled median value: 1.039 **A,B.** Dots represent median values. Gray shading represents 95% confidence intervals of the median. Dotted lines represent the median values observed for shuffled datasets (gray) and sampled datasets (red). X-axis labels represent the last common ancestor inferred for PRs and the corresponding estimated divergence times, in My. The numbers of gene pairs included in each groups are given in parentheses.

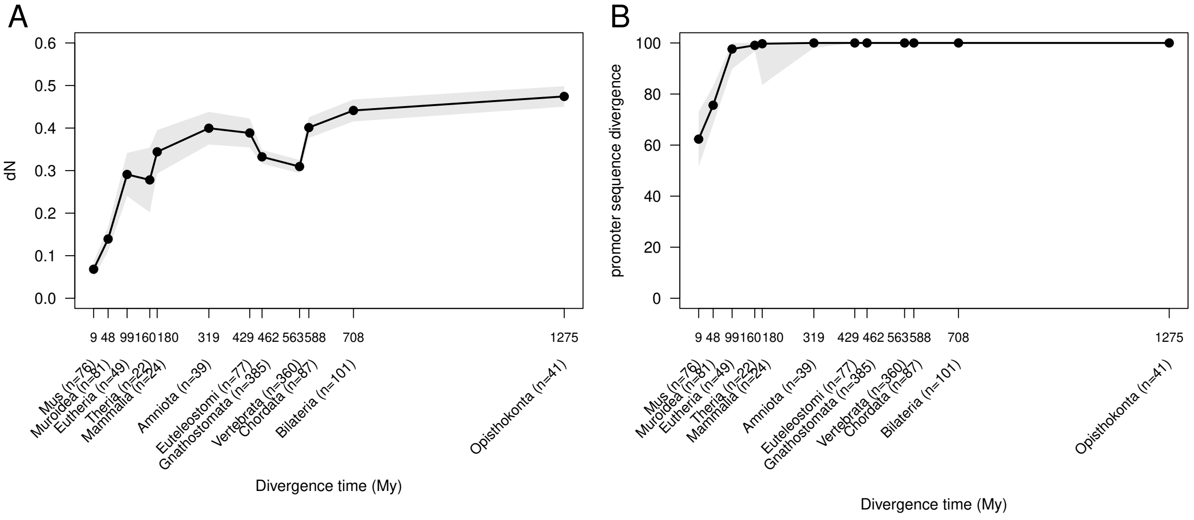


**Figure S24:** Dynamics of the protein sequence divergence and promoter sequence divergence with time. **A.** Distribution of the median rates of non-synonymous substitutions (dN) for paralogous gene pairs, as a function of their estimated divergence time. **B.** Distribution of the median promoter sequence divergence (computed as 100 - the percentage sequence identity, Materials and methods) for paralogous gene pairs, as a function of their estimated divergence time. **A,B.** Dots represent median values. Gray shading represents 95% confidence intervals. Dotted lines represent the median values observed for shuffled datasets (gray) and sampled datasets (red). X-axis labels represent the last common ancestor inferred for PRs and the corresponding estimated divergence times, in My. The numbers of gene pairs included in each group are given in parentheses.

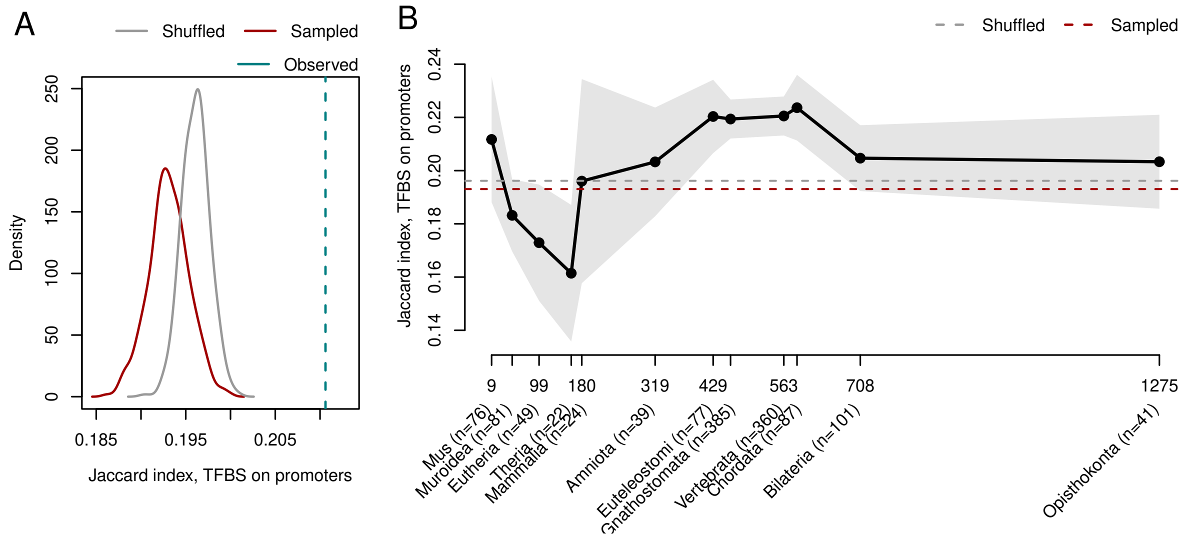


**Figure S25:** Similarity between the transcription factor binding sites (TFBS) identified on paralogous gene promoters. **A.** Density plot representing the distribution of the median TFBS promoter similarity (measured as the Jaccard index of the TFBS sets identified on two gene promoters, Materials and methods), for 1000 sampled datasets (red) and 1000 shuffled datasets (gray). The vertical bar represents the median TFBS promoter similarity of the paralogous gene pairs. **B.** Distribution of the median TFBS promoter similarity for paralogous gene pairs, as a function of their estimated divergence time. **A,B.** Dots represent median values. Gray shading represents 95% confidence intervals. Dotted lines represent the median values observed for shuffled datasets (gray) and sampled datasets (red). X-axis labels represent the last common ancestor inferred for PRs and the corresponding estimated divergence times, in My. The numbers of gene pairs included in each group are given in parentheses.

### Supplementary Tables

| Sample ID | Broad cell type or tissue | Detailed description | Cell line | Strain | Publication |
|---|---|---|---|---|---|
| preB_aged | B lymphocytes | Pre-B cells | - | C57BL/6 | Koohy et al. 2018 |
| preB_young | B lymphocytes | Pre-B cells | - | C57BL/6 | Koohy et al. 2018 |
| ESC | embryonic stem cells | Embryonic stem cells | J1 | 129S4/SvJae | Schoenfelder et al. 2015 |
| ESC_18 | embryonic stem cells | Embryonic stem cells | J1 | 129S4/SvJae | Schoenfelder et al. 2018 |
| ESC_wild | embryonic stem cells | Embryonic stem cells | E14Tg2a | 129P2/OlaHsd | Novo et al. 2018 |
| ESC_NKO | embryonic stem cells, Nanog KO | Embryonic stem cells, Nanog deficient | E14Tg2a | 129P2/OlaHsd | Novo et al. 2018 |
| EpiSC | epiblast stem cells | Epiblast stem cells | - | 129S2/SvPas | Novo et al. 2018 |
| ESd_starved | ES-derived hematopoietic progenitors | ES-derived multipotent hematopoietic progenitors | HPC-7 | 129S/SvEv-Gpi1 | Comoglio et al. 2018 |
| ESd_TPO | ES-derived hematopoietic progenitors | ES-derived multipotent hematopoietic progenitors | HPC-7 | 129S/SvEv-Gpi1 | Comoglio et al. 2018 |
| FLC | fetal liver | Fetal liver cells | - | C57BL/6 | Schoenfelder et al. 2015 |
| preadip_4H | pre-adipocytes | Pre-adipocytes | 3T3-L1 | Swiss albino | Siersback et al. 2017 |
| preadip_D0 | pre-adipocytes | Pre-adipocytes | 3T3-L1 | Swiss albino | Siersback et al. 2017 |
| preadip_D2 | pre-adipocytes | Pre-adipocytes | 3T3-L1 | Swiss albino | Siersback et al. 2017 |
| TSC | trophoblast stem cells | Trophoblast stem cells | TS-EGFP | 129/Sv and ICR | Schoenfelder et al. 2018 |

**Table S1:** List of PCHi-C samples used in this project (Laverré, Tannier, and Necsulea 2022).

| SRA Accession | PCHiC sample ID | Full description | Publication |
|---|---|---|---|
| SRR5826906 | ESd_starved | Subcellular RNA-seq Chromatin serum-starved Rep 1 | Comoglio et al. 2018 |
| SRR5826907 | ESd_starved | Subcellular RNA-seq Chromatin serum-starved Rep 2 | Comoglio et al. 2018 |
| SRR5826908 | ESd_TPO | Subcellular RNA-seq Chromatin TPO 30 min. Rep 1 | Comoglio et al. 2018 |
| SRR5826909 | ESd_TPO | Subcellular RNA-seq Chromatin TPO 30 min. Rep 2 | Comoglio et al. 2018 |
| SRR5826910 | ESd_starved | Subcellular RNA-seq Nucleoplasm serum-starved Rep 1 | Comoglio et al. 2018 |
| SRR5826911 | ESd_starved | Subcellular RNA-seq Nucleoplasm serum-starved Rep 2 | Comoglio et al. 2018 |
| SRR5826912 | ESd_TPO | Subcellular RNA-seq Nucleoplasm TPO 30 min. Rep 1 | Comoglio et al. 2018 |
| SRR5826913 | ESd_TPO | Subcellular RNA-seq Nucleoplasm TPO 30 min. Rep 2 | Comoglio et al. 2018 |
| SRR5826914 | ESd_starved | Subcellular RNA-seq Cytoplasm serum-starved Rep 1 | Comoglio et al. 2018 |
| SRR5826915 | ESd_starved | Subcellular RNA-seq Cytoplasm serum-starved Rep 2 | Comoglio et al. 2018 |
| SRR5826916 | ESd_TPO | Subcellular RNA-seq Cytoplasm TPO 30 min. Rep 1 | Comoglio et al. 2018 |
| SRR5826917 | ESd_TPO | Subcellular RNA-seq Cytoplasm TPO 30 min. Rep 2 | Comoglio et al. 2018 |
| SRR6512696 | preB_aged | Total RNA-seq preB aged 1 | Koohy et al. 2018 |
| SRR6512697 | preB_aged | Total RNA-seq preB aged 2 | Koohy et al. 2018 |
| SRR6512698 | preB_aged | Total RNA-seq preB aged 3 | Koohy et al. 2018 |
| SRR6512690 | preB_young | Total RNA-seq preB young 1 | Koohy et al. 2018 |
| SRR6512691 | preB_young | Total RNA-seq preB young 2 | Koohy et al. 2018 |
| SRR6512692 | preB_young | Total RNA-seq preB young 3 | Koohy et al. 2018 |
| SRR6766789 | ESC_wild | ESC_WT_1 | Novo et al. 2018 |
| SRR6766790 | ESC_wild | ESC_WT_2 | Novo et al. 2018 |
| SRR6766791 | ESC_wild | ESC_WT_3 | Novo et al. 2018 |
| SRR6766792 | ESC_NKO | ESC_NKO_1 | Novo et al. 2018 |
| SRR6766793 | ESC_NKO | ESC_NKO_2 | Novo et al. 2018 |
| SRR6766794 | ESC_NKO | ESC_NKO_3 | Novo et al. 2018 |
| SRR6766795 | EpiSC | EpiSC_1 | Novo et al. 2018 |
| SRR6766796 | EpiSC | EpiSC_2 | Novo et al. 2018 |
| SRR6766797 | EpiSC | EpiSC_3 | Novo et al. 2018 |
| SRR6766798 | EpiSC | EpiSC_4 | Novo et al. 2018 |
| SRR1605826 | ESC_18 | embryonic stem cells | Schoenfelder et al. 2018 |
| SRR1605832 | TSC | trophoblast stem cells | Schoenfelder et al. 2018 |
| SRR5297556 | preadip_4H | RNA_4h_Exp1 | Siersback et al. 2017 |
| SRR5297557 | preadip_4H | RNA_4h_Exp2 | Siersback et al. 2017 |
| SRR5297558 | preadip_D0 | RNA_D0_Exp1 | Siersback et al. 2017 |
| SRR5297559 | preadip_D0 | RNA_D0_Exp2 | Siersback et al. 2017 |
| SRR5297562 | preadip_D2 | RNA_D2_Exp1 | Siersback et al. 2017 |
| SRR5297563 | preadip_D2 | RNA_D2_Exp2 | Siersback et al. 2017 |

**Table S2:** List of RNA-seq samples used in this manuscript.

| GO Term | Description | P-value | FDR q-value | Enrichment |
|---|---|---|---|---|
| GO:0044238 | primary metabolic process | 5.64E-28 | 4.98E-24 | 2.22 |
| GO:0044237 | cellular metabolic process | 9.35E-28 | 4.13E-24 | 2.20 |
| GO:0006807 | nitrogen compound metabolic process | 4.19E-27 | 1.23E-23 | 2.31 |
| GO:0008152 | metabolic process | 4.83E-26 | 1.07E-22 | 2.03 |
| GO:0034660 | ncRNA metabolic process | 7.33E-26 | 1.3E-22 | 8.49 |
| GO:0071704 | organic substance metabolic process | 3.31E-24 | 4.88E-21 | 2.05 |
| GO:0034641 | cellular nitrogen compound metabolic process | 5.07E-24 | 6.4E-21 | 3.22 |
| GO:0090304 | nucleic acid metabolic process | 3.46E-23 | 3.82E-20 | 4.07 |
| GO:0046483 | heterocycle metabolic process | 1.13E-22 | 1.11E-19 | 3.37 |
| GO:0006139 | nucleobase-containing compound metabolic process | 6.84E-22 | 6.05E-19 | 3.13 |
| GO:0016070 | RNA metabolic process | 2.25E-21 | 1.81E-18 | 4.66 |
| GO:0006725 | cellular aromatic compound metabolic process | 1.31E-20 | 9.67E-18 | 3.23 |
| GO:0006399 | tRNA metabolic process | 2.8E-19 | 1.91E-16 | 11.84 |
| GO:0043170 | macromolecule metabolic process | 1.2E-18 | 7.57E-16 | 2.31 |
| GO:1901360 | organic cyclic compound metabolic process | 3.49E-18 | 2.05E-15 | 2.91 |
| GO:0034470 | ncRNA processing | 6.78E-17 | 3.75E-14 | 6.76 |
| GO:0016072 | rRNA metabolic process | 4.12E-16 | 2.14E-13 | 9.98 |
| GO:0044260 | cellular macromolecule metabolic process | 8.44E-16 | 4.15E-13 | 3.54 |
| GO:0006412 | translation | 1.88E-14 | 8.73E-12 | 10.89 |
| GO:0006364 | rRNA processing | 2.38E-14 | 1.05E-11 | 9.74 |

**Table S3:** GO enrichment results for old duplicated genes. The results were obtained with GOrilla, in the "single ranked list of genes" mode. Genes were ranked in decreasing order of their duplication age. In the results table, GO categories were ranked in order of increasing p-value. Only the first 20 GO categories are shown.

| GO Term | Description | P-value | FDR q-value | Enrichment |
|---|---|---|---|---|
| GO:0007608 | sensory perception of smell | 1.09E-31 | 9.67E-28 | 3.86 |
| GO:0007606 | sensory perception of chemical stimulus | 1.54E-31 | 6.8E-28 | 3.79 |
| GO:0007600 | sensory perception | 4.03E-15 | 1.19E-11 | 2.42 |
| GO:0006952 | defense response | 1.99E-13 | 4.39E-10 | 1.80 |
| GO:0007186 | G protein-coupled receptor signaling pathway | 9.12E-12 | 1.61E-8 | 1.68 |
| GO:0006955 | immune response | 1.53E-11 | 2.25E-8 | 1.87 |
| GO:0051707 | response to other organism | 2.83E-11 | 3.58E-8 | 2.57 |
| GO:0042742 | defense response to bacterium | 7.09E-11 | 7.83E-8 | 3.25 |
| GO:0098542 | defense response to other organism | 8.06E-11 | 7.92E-8 | 2.96 |
| GO:0009617 | response to bacterium | 6.58E-10 | 5.82E-7 | 2.51 |
| GO:0050877 | nervous system process | 1.05E-8 | 8.45E-6 | 1.88 |
| GO:0006959 | humoral immune response | 4.97E-8 | 3.66E-5 | 3.79 |
| GO:0051704 | multi-organism process | 6.65E-8 | 4.52E-5 | 1.83 |
| GO:0002376 | immune system process | 9.63E-8 | 6.08E-5 | 1.63 |
| GO:0043207 | response to external biotic stimulus | 1.6E-7 | 9.45E-5 | 2.06 |
| GO:0006954 | inflammatory response | 2.95E-7 | 1.63E-4 | 1.84 |
| GO:0003008 | system process | 3.1E-7 | 1.61E-4 | 1.52 |
| GO:0009607 | response to biotic stimulus | 5.02E-7 | 2.46E-4 | 2.00 |
| GO:0010951 | negative regulation of endopeptidase activity | 1.25E-6 | 5.84E-4 | 16.57 |
| GO:0048002 | antigen processing& presentation of peptide antigen | 1.67E-6 | 7.38E-4 | 31.66 |

**Table S4:** GO enrichment results for young duplicated genes. The results were obtained with GOrilla, in the "single ranked list of genes" mode. Genes were ranked in increasing order of their duplication age. In the results table, GO categories were ranked in order of increasing p-value. Only the first 20 GO categories are shown.
